# Convergent evolution of somatic escape variants in *SERPINA1* in the liver in alpha-1 anti-trypsin deficiency

**DOI:** 10.1101/2024.09.02.610686

**Authors:** Natalia Brzozowska, Lily YD Wu, Vera Khodzhaeva, William J Griffiths, Adam Duckworth, Hyunchul Jung, Tim H H Coorens, Yvette Hooks, Joseph E Chambers, Peter J Campbell, Stefan J Marciniak, Matthew Hoare

## Abstract

Somatic variants accumulate in non-malignant tissues with age^1,2^. Functional variants leading to clonal advantage of hepatocytes accumulate in the liver from patients with acquired chronic liver disease (CLD)^3–5^. Whether these somatic variants are common to CLD from differing aetiologies is unknown. We analysed somatic variants in the liver from patients with genetic CLD from alpha-1 anti-trypsin (A1AT) deficiency or haemochromatosis. We show that somatic variants in *SERPINA1*, the gene encoding A1AT, are strongly selected for in A1AT deficiency, with evidence of convergent evolution. Acquired variants cluster at the 3’ end of *SERPINA1* leading to C-terminal missense or truncation variants of A1AT. *In vitro,* C-terminal truncation variants abrogate disease-associated A1AT polymerisation and retention in the endoplasmic reticulum, supporting the C-terminal domain swap mechanism. Therefore, somatic escape variants from a deleterious germline variant are selected for in A1AT deficiency, suggesting adaptive functional somatic variants are disease-specific in CLD and point to disease-associated mechanisms.

---

Chronic liver disease (CLD) leading to cirrhosis accounts for 1 in 25 deaths globally^6^. The aetiology of CLD is changing with fewer cases attributable to viral hepatitis, but an increased incidence of obesity and type 2 diabetes driving a worldwide increasing prevalence of metabolic-dysfunction associated steatotic liver disease (MASLD). Beyond these, significant numbers of CLD cases relate to germline genetic disorders, including haemochromatosis and alpha-1 anti-trypsin (A1AT) deficiency^7^. Haemochromatosis, common in patients in northern Europe, most frequently results from inherited mutations within *HFE* or other genes involved in iron absorption and metabolism^8^, leading to deposition of hepatocellular iron and subsequent cellular stress. A1AT deficiency results from inherited homozygous K366E mutations of the *SERPINA1* gene, encoding the Z-variant of the A1AT protein^9^. Unlike the wild-type M-A1AT, which is secreted into the serum to inhibit neutrophil elastase, the Z-variant polymerises in the hepatocyte endoplasmic reticulum (ER) leading to ER stress and cell death. There are currently no treatment options for patients with progressive PiZZ A1AT deficiency, other than liver transplantation.

Somatic acquisition of functionally advantageous variants with subsequent clonal expansion points to the differential selection pressure that operates within the diseased microenvironment. In our previous work on MASLD and alcohol-related liver disease, we identified convergent evolution of functional variants in metabolism genes, including *FOXO1, CIDEB* and *GPAM*, that drive clonal hepatocyte expansion, likely through modulation of carbohydrate and lipid metabolism^4^, known to be dysregulated in MASLD. Loss-of-function in the homologous genes lead to clonal expansion of hepatocytes in mouse models of MASLD^10^; common germline variants of *GPAM*^11,12^ or rare predicted loss-of-function germline variants of *CIDEB*^13^ protect against the development of MASLD. However, it is likely that different diseases leading to specific microenvironmental stressors will drive differential selective advantage of driver variants in similar contexts: liver tissues from patients in the southern United States, predominantly with viral hepatitis, develop a different spectrum of somatic variants^3^. Therefore, identification of advantageous somatic variants that arise in specific disease contexts potentially points to underlying functional mechanisms; mimicry would be a rational therapeutic strategy, especially in liver diseases where no therapeutics currently exist.

In this study we explored somatic variants and clonal dynamics in the liver of patients with end-stage CLD caused by haemochromatosis and A1AT deficiency. In a group of patients with similar background demographics to our previous cohort with MASLD^4,5^, but with CLD of distinct aetiology, we hypothesised that disease-specific pressure would drive expansion of clones containing distinct functional variants in genetic CLD: both diseases have pathological mechanisms that do not overlap with each other or with steatotic liver disease (SLD). We obtained explanted liver tissue from 5 patients with homozygous PiZZ-A1AT deficiency (Supplementary Table 1, supplementary figure 1) and 5 patients with haemochromatosis; all had significant hepatocellular iron deposition and included one patient with homozygous C282Y mutations, one with compound C282Y / H63D mutations and three with haemochromatosis of uncertain cause (Supplementary Table 2) without risk factors for secondary haemochromatosis. All were undergoing liver transplantation for liver failure, hepatocellular carcinoma (HCC) or both. To identify somatic variants, we initially performed whole-genome sequencing (WGS) in 1 patient with A1AT-deficiency and 4 with haemochromatosis, but thereafter used only whole exome sequencing (WES) in the remaining cases, to focus on coding variation. We performed laser-capture microdissection and WGS / WES (Fig. 1A) of 305 laser microdissections of liver tissue achieving a mean coverage of 42X (Fig. 1B) for both sequencing platforms. To consider exome and genome datasets together, we reported the number of exonic coding mutations (Fig. 1B). There was considerable heterogeneity in variant burden both between and within patients (Supplementary Fig. 2A), but no significant differences between A1AT deficiency, haemochromatosis or previously sequenced liver samples from patients with SLD^4^ and no differential representation of mutational signatures (Supplementary Fig. 2B). Copy number changes and structural variants (called in whole-genome samples) occurred in moderate numbers across haemochromatosis patients and the one A1AT deficiency patient with WGS data (Supplementary Fig. 2A). One clone demonstrating chromothripsis involving chromosomes 7 and 10 was detected in the liver of a patient with haemochromatosis undergoing transplantation for liver failure without HCC (Supplementary Fig. 2C), accounting for 30/48 of SVs and 30/34 CNVs in that donor.

**Figure 1.**
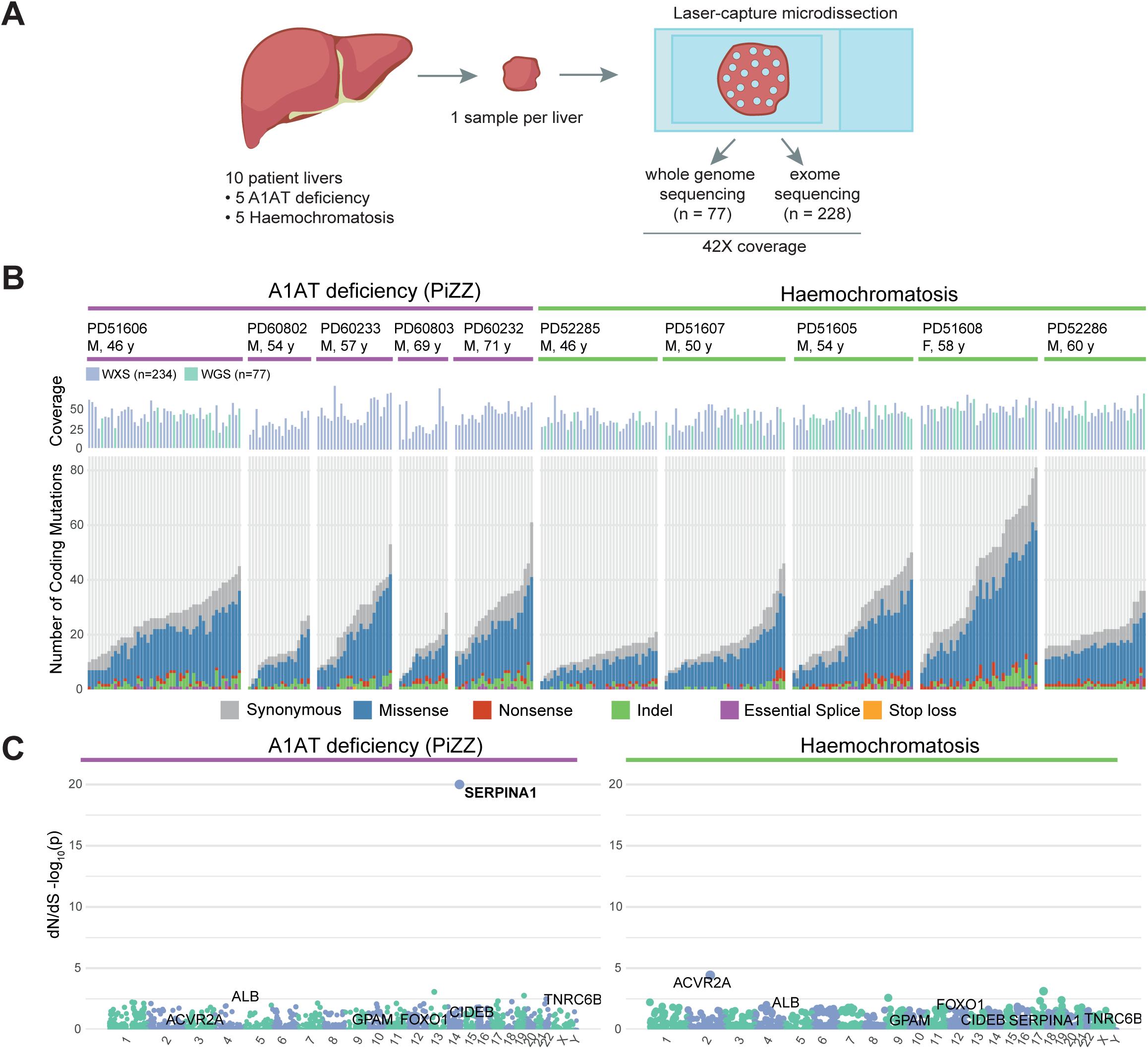
Liver somatic variants in alpha-1 anti-trypsin (A1AT) deficiency and haemochromatosis. (A) Overview of the hierarchical experimental design – 10 livers from patients with A1AT deficiency and haemochromatosis were sampled. These samples underwent laser capture microdissection to generate 305 individual microdissections; isolated DNA was then sequenced using whole genome or whole exome approaches. (B) Top panel: mean sequencing coverage in each microdissection. Donors are ordered by disease and age. The number of single nucleotide variants and indels detected per microdissection within the coding exome across whole genome (n = 77) and exome (n = 228) sequencing data. Variants are coloured by type. (C) Genes under positive selection in A1AT deficiency and haemochromatosis cohorts: Manhattan plots showing the distribution of p-values testing for gene-level non-synonymous variants in A1AT deficiency (left panel) and haemochromatosis (right panel). Labelled are genes found to be under positive selection in A1AT deficiency, haemochromatosis or, previously, in SLD. Genes are ordered by genomic position, and those with significant q values (<0.1) are highlighted in bold.

We analysed which genes were recurrently affected by acquired non-synonymous variants more than expected by chance^14^: across all protein-coding genes, we found that acquired variants in *SERPINA1* were under strong positive selection (q < 2×10^-16^) (Fig. 1C) in patients with A1AT deficiency, but not patients with haemochromatosis (q = 1) or SLD^4,5^ (q = 0.73). We did not identify any enrichment for somatic variants in any gene previously associated with iron metabolism^8^, including *HFE*, in either cohort. Further, no acquired variants were positively selected for in the patients with haemochromatosis; this may reflect a lack of statistical power given the small numbers of patients examined. Genes previously found to be under positive selection in liver tissue from patients with SLD^4^ or liver cancer^15^ did not appear to be under positive selection in liver tissue from either the A1AT deficiency or haemochromatosis cohorts: *CIDEB* dN/dS ratio 72.1 (95% CI 4.0 - 367.9; q = 1); *FOXO1* ratio 0.0 (0.0 – 43.4; q = 1); *GPAM* ratio 0.0 (0.0 – 40.6; q = 1). Therefore, liver diseases of differing aetiology drive the expansion of hepatocyte clones with distinct somatic variants.

In the five patients with A1AT deficiency, we observed 17 indels, 2 missense and 3 nonsense mutations in *SERPINA1*. All five patients had genetic evidence of convergent evolution with multiple independent clones containing distinct *SERPINA1* variants within the same piece of randomly-selected liver tissue (Fig. 2). One patient had 11 independent clones with *SERPINA1* variants within 3cm^2^ of liver tissue, with one clone containing two frameshift deletions in the gene (Fig. 2A,B). The further four patients had two or three clones with independent *SERPINA1* variants (Fig. 2C); most non-synonymous mutations in *SERPINA1* in the A1AT deficiency cohort had variant allele fractions (VAFs) between 0.2 and 0.4, well within the VAF distribution for passenger mutations in the respective samples, indicative of being heterozygous, rather than homozygous, in the major clone (Supplementary Fig. 3). Therefore, heterozygous somatic variants in *SERPINA1* provide selective advantage in A1AT deficiency, which only develops in patients with homozygous germline Z variants of *SERPINA1*.

**Figure 2.**
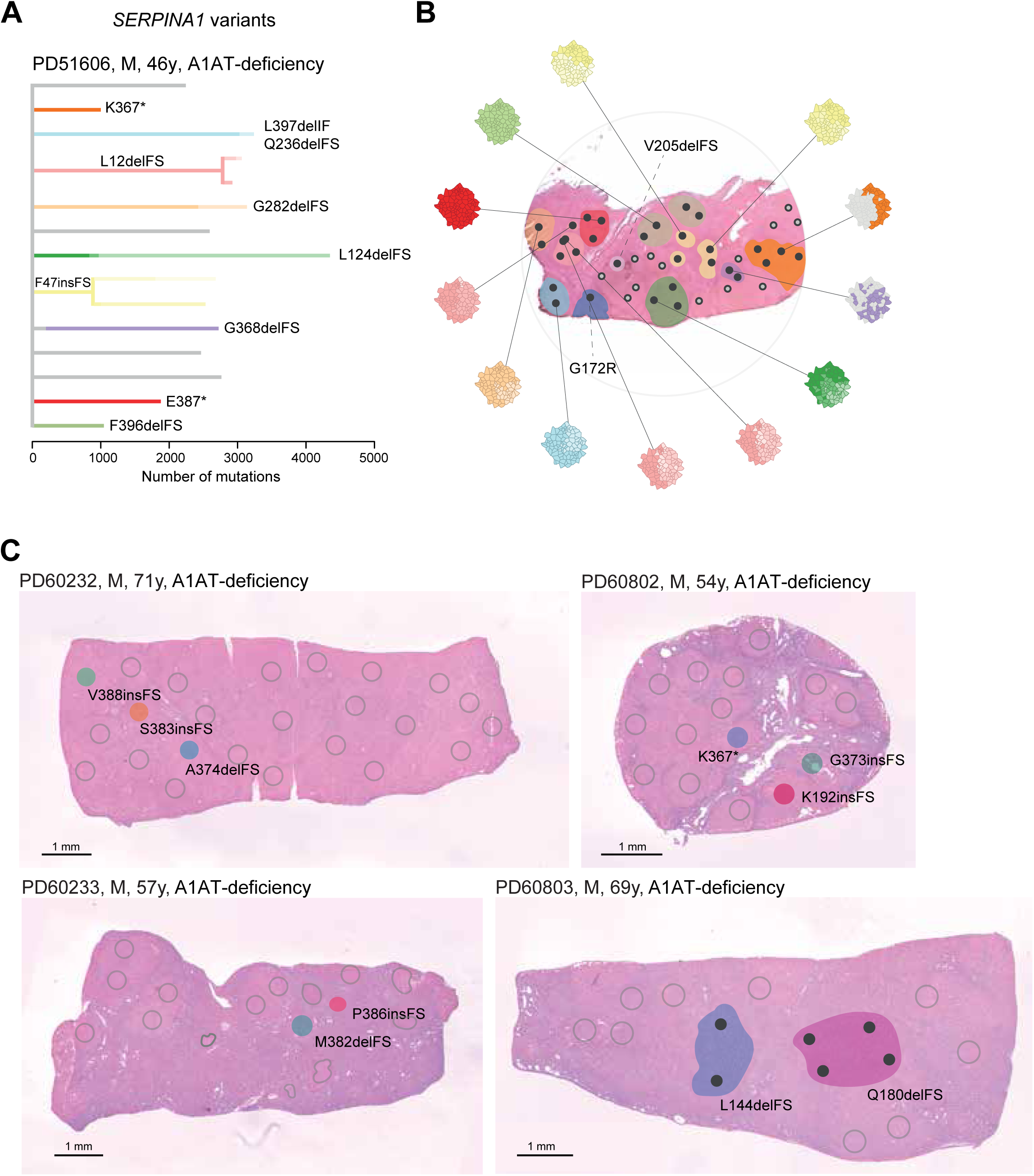
Convergent evolution of somatic variants of *SERPINA1* in alpha-1 anti-trypsin (A1AT) deficiency. (A) Phylogenetic tree of clonal structure from liver sample PD51606, with coloured branches showing independently acquired *SERPINA1* mutations. Solid lines indicate that nesting is in accordance with the pigeonhole principle; dashed lines indicate that nesting is in accordance with the pigeonhole principle, assuming that hepatocytes represent <100% of cells. (B) Spatial mapping of the clones from the phylogenetic tree onto an H&E-stained photomicrograph of the patient’s liver biopsy, with *SERPINA1*-mutant clones coloured to match the tree. Each dot signifies a microdissection, with solid black dots indicating LCM microdissections harbouring a *SERPINA1* mutation. For WGS samples, clonal cell fraction is depicted, with a fully filled disk signifying the entire LCM sample and *SERPINA1* mutant cell fraction depicted by the degree of disk filling. Dashed lines indicate *SERPINA1* mutations identified exclusively in WES samples. Nodules are colour-highlighted where all microdissections within the nodule contain the same *SERPINA1* mutant clone. (C) Spatial location of *SERPINA1*-mutant clones in the H&E-stained liver of the remaining 4 A1AT deficiency patients (coloured area), in addition to *SERPINA1* wild-type clones (grey circles). Scale bars 1mm.

We were interested in exploring the potential protein coding effects of the somatic variants in *SERPINA1*. Notably, the identified variants in the A1AT deficiency cohort showed strong clustering of truncating variants within the last exon of *SERPINA1* (Fig. 3A), not seen in the previous SLD cohort (Supplementary Fig. 4): amongst 1590 genomes from these 35 SLD patients, three patients had a heterozygous PiMZ *SERPINA1* genotype with only one identified frame-shift indel at S6 of *SERPINA1* in these liver samples. In the A1AT deficiency cohort, all frameshift indels and nonsense mutations in the last exon are predicted to result in the loss of at least 19 amino acids from the C-terminus of the encoded protein, in addition to partial modification of amino acid sequence in the case of frameshift indels (Fig. 3B). The recurrence and convergent evolution of similar variants suggested that acquired C-terminal missense or truncation *SERPINA1* variants provide a selective advantage to hepatocytes in A1AT deficiency through common functional mechanisms.

**Figure 3.**
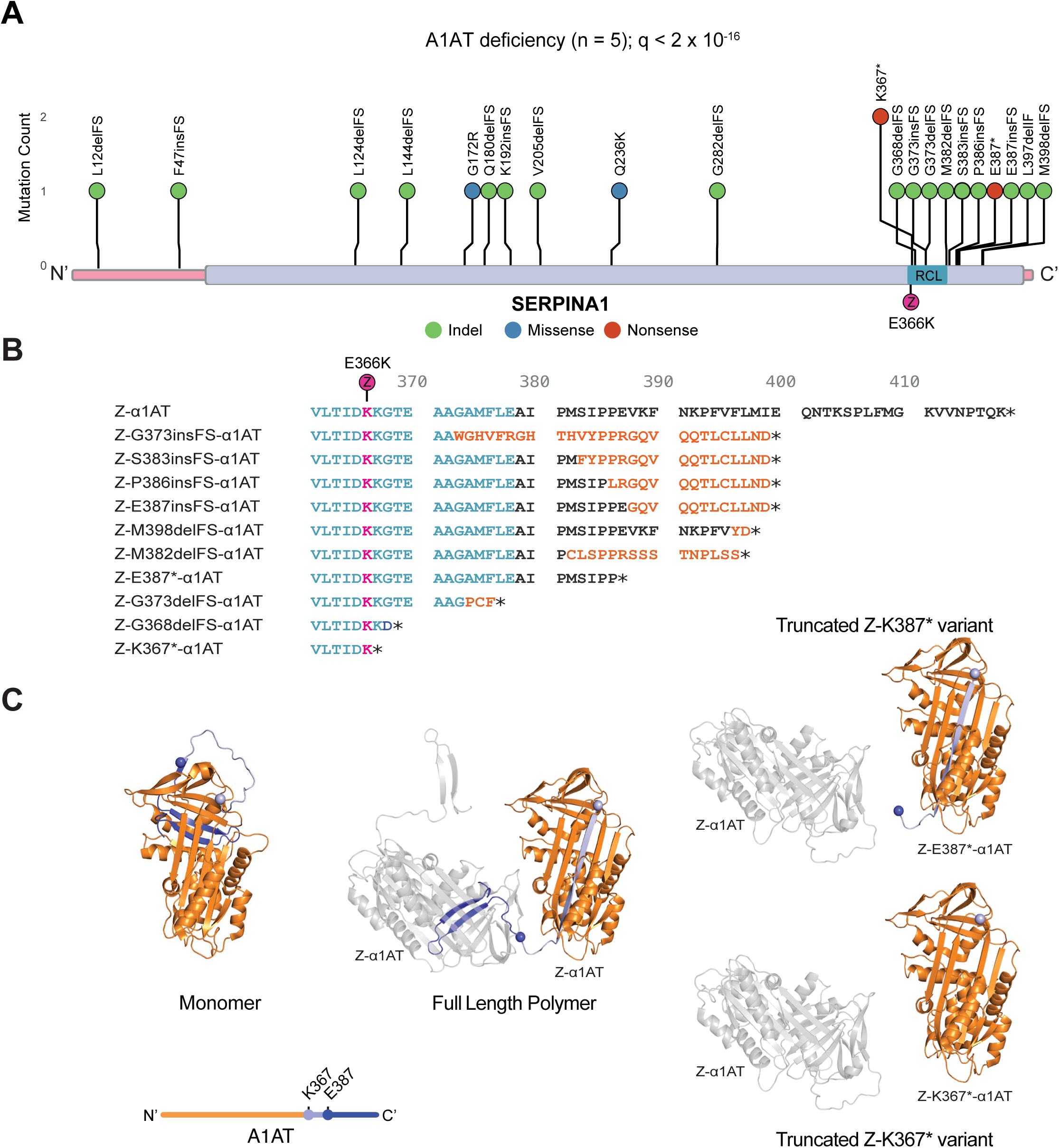
Recurrent variants of *SERPINA1* are predicted to lead to C-terminal protein truncations. (A) Distribution of non-synonymous variants detected in *SERPINA1* in liver samples from patients with A1AT deficiency (n = 5). Coding sequence of the gene represented on the x-axis, with exons in pink and protein domains in purple. Position of the germline PiZ variant (E366K) shown in pink; RCL, reactive centre loop. (B) Effect of C-terminal variants on Z-A1AT protein sequence; sequence alignment of Z-A1AT amino acids 361-418. Reactive centre loop is highlighted in blue; amino acid sequence modified by frameshift in orange; E366K (PiZ) variant indicated in pink circle; stop codon denoted by *. (C) Left: Cartoon representing the crystal structure of native A1AT derived from Protein Data Bank (PDB):1QLP^17^. The position of residues K367 (magenta) and E387 (blue) are shown by coloured spheres, with downstream residues coloured similarly. A linear schematic of the protein shows relative mutation positioning using the same colour code (upper). Middle: The crystal structure of two protomers in a Z-A1AT polymer from PDB:3T1P^18^ showing insertion of donor protomer residues downstream of K367 (magenta sphere) into the acceptor protomer. Right: A structural model of two protomers in a polymer (PDB:3T1P) highlighting the incompatibility of Z-K367* and Z-E387* C-terminal donation to a polymer.

Somatic variants leading to truncation (Fig. 3C, right) or sequence changes (Supplementary Fig. 5) in the C-terminus of A1AT have the potential to disrupt native folding of A1AT. However, *in silico* modelling predicts that this should not occur. The recurrent *SERPINA1* variants affect the A1AT domain including the reactive centre loop (RCL) and downstream C-terminus, which have previously been shown to mediate homopolymerization of Z-A1AT (Fig. 3C)^16^. In its native fold, the C-terminal 26 amino acids of A1AT form a β-hairpin that inserts into, and completes β-sheet B, with β-sheet A adopting a 5-stranded conformation^17^ (Fig. 3C, middle). Polymers arise through the completion of β-sheet B in an “acceptor” protomer by the C-terminal β hairpin of a second “donor” protomer^16,18^. This causes the acceptor molecule to adopt a hyper-stable, 6-stranded β-sheet - a conformation that translocates its own orphaned C-terminus to the opposite pole of the molecule, where it can act as a donor to drive polymer extension (Fig. 3C, middle). Therefore, somatic variants that impose truncation or sequence changes in the C-terminus of A1AT have the potential to disrupt polymerisation of the Z protein through lost or reduced ability (Supplementary Fig. 5C-E) to donate the C-terminus into neighbouring Z-protomers, but should not affect their ability to act as “acceptors”.

To explore the potential mechanisms, including polymerisation, underlying the selective advantage of hepatocytes containing C-terminal truncation variants of Z-A1AT we modelled these *in vitro*: we chose to study two variants with acquired premature stop codons: Z-K367* and Z-E387* (Fig. 3B,C). To study cellular localisation of Z-A1AT with these variants, we conducted live-cell imaging of HaloTag-fused A1AT in an established CHO cell model that lacks endogenous A1AT expression^19,20^. Both M- and Z-A1AT HaloTag fusion proteins colocalised with the ER marker protein moxGFP-KDEL, whilst only the M form also localised to perinuclear structures resembling the Golgi apparatus. Expression of both M- and Z-forms of A1AT can lead to fragmentation of the ER network giving rise to punctate structures termed ER-inclusions (Fig 4A). We quantified ER inclusion formation in cells expressing HaloTagged M, Z, Z-K367* or Z-E387* variants. Z-A1AT showed a greater propensity to form ER inclusions than M-A1AT (Fig. 4A,B), as reported previously^20,21^; by contrast, Z-K367* or Z-E387* led to a similar propensity to form ER inclusions as seen with wild-type M-A1AT expression (Fig. 4A,B). Therefore, the C-terminal truncation variants of Z-A1AT show reduced accumulation within, and disruption of, the ER.

**Figure 4.**
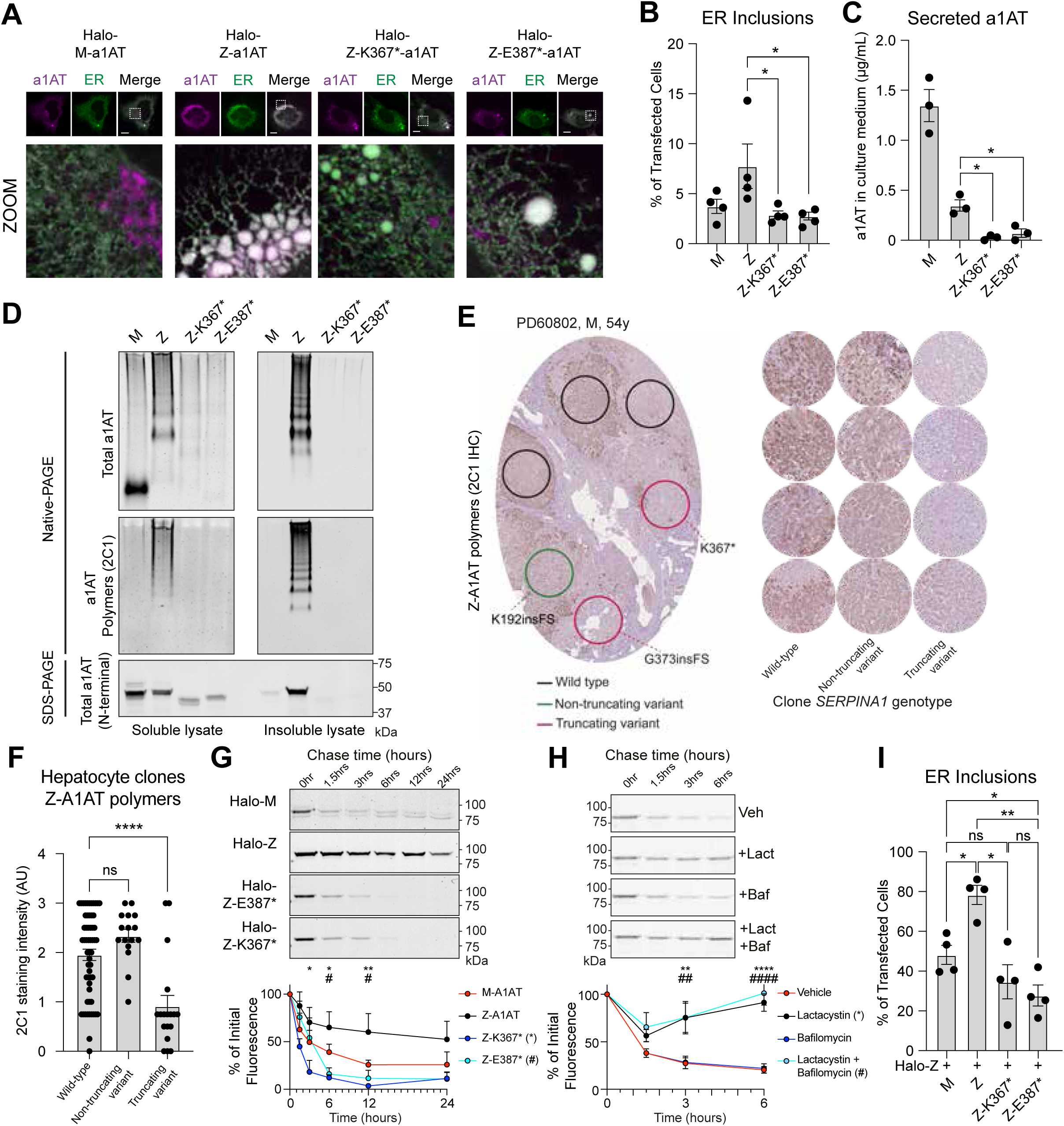
Somatic C-terminal truncation variants of *SERPINA1* inhibit polymerisation of PiZ alpha-1 anti-trypsin and ER disruption. (A) Airyscan fluorescence micrographs of CHO cells acquired 48 hours after transient transfection with an expression vector encoding HaloTagged variants of A1AT (M, Z, Z-K367*, and Z-E387*, in magenta) with the ER-marker protein (moxGFP-KDEL, in green). Dashed line boxes on merged channel images denote area expanded in zoom images. Scale bar 10μm. (B) CHO cells prepared as in (A) were analysed by confocal microscopy to establish the proportion of transfected cells displaying regions containing ER-inclusions (characterised by a region of fragmented tubular ER network); Analysis by one-way ANOVA with Sidak’s correction for multiple hypothesis testing, * P ≤ 0.05. (C) Sandwich ELISA of conditioned culture medium from CHO cells transiently transfected to express A1AT variants (M, Z, Z-K367*, and Z-E387*). Capture was performed with a polyclonal antibody raised against M antitrypsin and detection performed using the non-conformationally selective mAb_3C11_ antibody; Columns represent means ± SEM; * P ≤ 0.05. (D) CHO cells were transfected to express untagged A1AT variants 48 hours prior to SDS- and Native-PAGE of soluble and insoluble fractions of cell lysates from an equivalent cell mass. Native-PAGE immunoblotting was performed using antibodies recognising the total (a0409) and polymerised (mAB_2C1_) A1AT pools. SDS-PAGE immunoblotting was performed using an antibody raised against an N-terminal epitope of A1AT present in all variants (MA5-15521). (E) Left panel: Exemplar photomicrograph of liver tissue stained for Z-A1AT polymers by 2C1 immunohistochemistry from sample PD60802 showing the staining pattern across the liver sample; circles indicate regions with microdissected hepatocyte clones analysed for WES/WGS (as in Figure 1) in adjacent section and demonstrated to have wild-type or acquired variants in *SERPINA1*; right panel: exemplar photomicrographs of 2C1 immunohistochemistry in liver tissue from regions with wild-type or acquired variants in *SERPINA1* across the A1AT-deficiency cohort. (F) Quantification of 2C1 immunohistochemistry (as in (E)) staining intensity on liver sections from 5 patients with A1AT-deficiency scored (0-3); Analysis by one-way ANOVA with Sidak’s correction for multiple hypothesis testing, **** P ≤ 0.0001. (G) SDS-PAGE gels of CHO cell lysates expressing HaloTagged A1AT variants pulse-labelled with JF646 HaloTag ligand, then harvested at the indicated timepoints post-labelling. Quantitation of JF646 labelled HaloTag-A1AT protein bands is shown. Analysis by two-way ANOVA with Sidak’s correction for multiple hypothesis testing, * and # denote significance of Z vs Z-K367* or Z vs Z-E387* respectively. (H) Pulse labelling of HaloTagged Z-K367* performed as in (G), in the presence of DMSO (Vehicle), Lactacystin (proteosome inhibitor, 5µM), Bafilomycin (lysosomal inhibitor, 100nM), or both. Analysis by two-way ANOVA with Sidak’s correction for multiple hypothesis testing, * and # denote significance of Vehicle vs Lact or Vehicle vs both inhibitors respectively. (I) Proportion of CHO cells displaying ER inclusions when co-expressing HaloTagged Z-A1AT with untagged variants of A1AT; Analysis by one-way ANOVA with Sidak’s correction for multiple hypothesis testing, * P ≤ 0.05, ** P ≤ 0.01.

The localisation of the M-variant apparently to the Golgi apparatus supports the expected secretory trafficking of wild-type HaloTag-M-A1AT, with ER retention of the Z-variant^22^ (Fig. 4A). Despite reduced ER retention, the HaloTagged Z-K367* and Z-E387* forms of A1AT showed no evidence of secretory trafficking: quantitative sandwich ELISA (Fig. 4C) and immunoblotting using an N-terminal directed antibody (Supplementary Fig. 6A) for secreted (untagged) A1AT in conditioned media from transfected CHO cell cultures indicated that cells expressing Z-K367* and Z-E387*, secrete very little A1AT protein. Therefore, the C-terminal truncation variants of Z-A1AT do not accumulate in the ER, but do not restore secretion of A1AT.

We next investigated ER handling of C-terminal truncation variants, by assessing their mobility within the ER lumen using single particle tracking (Supplementary Fig. 6B). HaloTagged A1AT variants were expressed in COS7 cells, labelled with a far-red photoactivatable fluorescent ligand^23^, and single molecule velocities calculated using a simple linear assignment problem algorithm^24^. As shown previously, single Z-A1AT particles displayed a significant reduction in velocity compared to M-A1AT (Supplementary Fig. 6B)^20^. Z-K367* and Z-E387* also showed significant reduced velocity, suggesting these C-terminal truncations do not restore normal ER handling. Reduced mobility of Z-K367* and Z-E387* could indicate either an increase in protein hydrodynamic volume via polymerisation or interactions with ER quality-control factors. Therefore, C-terminal truncation variants of A1AT had reduced propensity to form ER inclusions, but these variants failed to rescue impaired molecular mobility within the ER or trafficking through the secretory pathway.

Based on the predicted protein sequences, we hypothesised that the C-terminal truncation variants could impair aberrant polymerisation of Z-A1AT. We assessed A1AT polymer formation in CHO cells expressing untagged A1AT variants by polyacrylamide gel electrophoresis under native non-denaturing (Native-PAGE), and denaturing (SDS-PAGE) conditions, followed by immunoblotting. As Z-A1AT polymers partition between soluble and insoluble pools within the ER^7,20,25,26^, insoluble proteins were first separated from lysates by centrifugation and resuspended in their initial volume to assess stoichiometry with soluble protein. Native-PAGE immunoblotting of the soluble A1AT pool showed that M-A1AT migrated predominantly as a species of low molecular weight representing folded monomers (Fig. 4D, upper left panel). Z-A1AT separated as a ladder of higher molecular weight species in soluble and insoluble pools, that were confirmed as A1AT polymers by the conformation-specific antibody mAb_2C1_ (Fig. 4D, middle panels). The C-terminal truncation variants Z-K367* and Z-E387* lack β-strands 4B and 5B required for C-terminal donation in polymerisation, and accordingly showed low abundance of higher molecular weight species (Fig. 4D middle left panel) that react with polymer-specific mAb_2C1_ (Fig. 4D, middle right panel) despite containing the germline E366K Z-variant. Immunoblotting of lysates separated by SDS-PAGE confirmed lower expression of C-terminal truncation variants compared to Z-A1AT, but that they predominantly partition to the soluble fraction of lysates, in contrast to Z-A1AT (Fig. 4D, lower panel). Therefore, acquired C-terminal truncation variants of Z-A1AT have reduced ability to polymerise compared to germline Z-variants of A1AT *in vitro*.

We next explored whether hepatocyte clones that contained *SERPINA1* variants had evidence of reduced polymer formation *in vivo*. We used immunohistochemistry with the Z-A1AT polymer-specific 2C1 antibody on serial sections from the 5 liver samples in the A1AT-deficiency cohort, where we had already conducted spatial DNA-sequencing on the adjacent section (Fig 4E). We then defined areas of hepatocytes with wild-type *SERPINA1*, non-truncating variants of *SERPINA1*, or C-terminal truncation variants before assessing the intensity of 2C1 immunostaining in those areas. Hepatocytes with non-truncating *SERPINA1* variants had similar polymer burden to areas with wild-type *SERPINA1,* whereas hepatocytes with C-terminal truncation variants of *SERPINA1* had significantly lower polymer burden than wild-type areas (Fig. 4E,F). Therefore, C-terminal truncation and missense variants of Z-A1AT have reduced ability to polymerise *in vivo* compared to germline Z-variants of A1AT. These findings also raise the interesting question of whether proximal non-truncating variants are truly advantageous or whether these are simply passenger mutations.

As Z-K367* and Z-E387* variants were heterozygous (VAF ≤0.5) with native Z-A1AT expressed from the other allele in hepatocyte clones (Supplementary Fig. 3), we explored whether the reduction in A1AT accumulation was due to their potential to behave as “acceptor” protomers in forming mixed polymers with Z-A1AT, but preventing onward polymerisation through an inability to “donate” to the next Z monomer (Fig. 3C). HaloTagged A1AT variants were constitutively expressed in CHO cells that also conditionally-express untagged Z-A1AT under doxycycline (dox) control, before affinity purification of the Halotagged A1AT species (Supplementary Fig. 6C). SDS-PAGE immunoblotting of the input samples showed expression of the higher molecular-weight HaloTagged A1AT variants, in addition to the lower molecular weight untagged Z-A1AT. Affinity purification of HALO and then immunoblotting showed that the HALO-tagged Z-A1AT pulled-down lower molecular weight untagged Z-A1AT; similarly, purification of both Z-K367* and Z-E387* variants retained the ability to pull-down untagged Z-A1AT consistent with their ability to bind to native Z-A1AT derived from the unaffected allele. Therefore, whilst the C-terminal truncation variants have reduced ability to polymerise, they retain the ability to bind to Z-A1AT acting as “acceptor” protomers, but presumably preventing onward polymerisation due to defective “donation”.

It remained possible that reduced ER disruption could reflect impaired polymerisation or simply lower expression levels of the C-terminal Z-A1AT variants compared to the germline M- and Z-A1AT variants (Fig. 4D, lower panel). We explored this using a pulse-chase strategy, treating the cells with Janelia-Fluor reporters that covalently bind to HaloTagged A1AT variants in CHO cells. Consistent with previous data, we found that Z-A1AT had a longer half-life than M-A1AT (Fig. 4G). However, we found that both Z-K367* and Z-E387* variants had a significantly shorter half-life than Z-A1AT, similar to M-A1AT. To probe how these C-terminal variants of A1AT undergo degradation we utilised small molecule inhibitors of the two major protein degradation pathways: lactacystin as an inhibitor of the proteosome and bafilomycin A1 as an inhibitor of the lysosomal function. Whereas lysosomal inhibition had no effect upon the half-life of Z-K367*, proteosomal inhibition completely prevented the degradation of Z-K367* (Fig. 4H). Therefore, lower expression of the C-terminal truncation Z-A1AT variants compared to germline Z-A1AT is due to proteasomal degradation leading to a shorter half-life.

To understand whether Z-A1AT truncation variants can lead to reduced ER inclusions and disruption due to a loss-of-function from lower expression or gain-of-function, preventing polymerisation in a dominant-negative fashion, we transduced CHO cells with bicistronic vectors containing Z-A1AT, in addition to M-A1AT, Z-A1AT, Z-K367* or Z-E387*. As expected, CHO cells co-expressing Z-A1AT / Z-A1AT had significantly more inclusions than cells co-expressing Z-A1AT / M-A1AT (Fig. 4I). Cells expressing Z-K367* with Z-A1AT had similar levels of inclusions to Z-A1AT / M-A1AT-expressing cells. However, CHO cells with Z-E387* with Z-A1AT had significantly fewer inclusions than Z-A1AT / M-A1AT-expressing cells suggesting that these variants certainly have reduced polymerisation from reduced protein expression, but might also be able to dominantly inhibit Z-A1AT polymerisation. This will require further exploration, given that a dominant-negative function would have significant therapeutic implications.

Overall, we have identified convergent evolution of somatic variants in *SERPINA1* in the liver of patients with end-stage A1AT deficiency. The liver of each patient with PiZZ A1AT deficiency contained multiple hepatocyte clones with independent acquisition of variants of *SERPINA1*, the gene whose germline variant protein drives the disease process. Remarkably, most of these acquired variants lead to missense or truncation variants of the C-terminus of the A1AT protein and support the idea that *SERPINA1* variants engender cellular advantage by mitigating Z-A1AT polymerization^16,27^ and subsequent cellular stress. *In vitro* modelling of two of the C-terminal truncating variants has shown that these are associated with reduced Z-A1AT polymerisation as well as reduced accumulation within and disruption of the endoplasmic reticulum.

Avoidance of A1AT polymerisation and subsequent endoplasmic reticulum dysfunction would provide significant selective advantage for hepatocyte clones in A1AT deficiency compared to neighbouring Z-A1AT hepatocytes and therefore drive clonal expansion (Supplementary Fig. 7). This has important therapeutic implications in a disease that currently has no available therapies, beyond transplantation. Although therapies based on liver-directed RNA-interference, suppressing the expression of Z-A1AT, have shown promise in mouse models^28^ and preliminary human trials^29^, targeted blockade or deletion of the C-terminus of A1AT may represent an alternative strategy for suppression of polymer formation in A1AT deficiency. All of these strategies would promote hepatocyte survival and prevent progressive liver disease, but none would restore normal A1AT secretion or serum levels and therefore would likely not prevent progressive lung disease due to A1AT deficiency.

The effect of disease-associated germline variants upon subsequent somatic mutagenesis is relatively unexplored outside of hereditary cancer syndromes^30,31^. There are rare descriptions of acquired somatic revertants that correct germline variants in deleterious germline disorders: somatic genetic rescue (SGR) has been seen in severe combined immunodeficiency (SCID), where an individual with inherited biallelic *ADA* mutations acquired a reversion mutation, leading to outgrowth of a healthy lymphocyte population^32^. SGR has been described in muscle tissue of patients with inherited Duchenne muscular dystrophy^33,34^, as well as patients with hereditary tyrosinaemia where somatic revertants were identified in explanted liver tissue and their extent correlated with clinical severity^35^. Although similar to previous SGR, our identified variants in A1AT deficiency are not true revertants, but escape variants at a distinct site in the gene, mitigating the effect of a deleterious germline variant. Somatic escape variants in distinct genes have previously been identified in some cases of Shwachman-Diamond syndrome where acquired variants in *EIF6* act to mitigate the deleterious effects of germline EFL1 or SBDS mutations^36^.

Combined with our previous data in SLD, where distinct functional variants in metabolism genes were identified, these data suggest that differing disease processes will drive expansion of clones containing disease-specific variants providing selective advantage in the face of specific microenvironmental pressures. Identification of the advantageous variants in different tissues, populations and disease states has the potential to reveal functional gene targets, but also reveal specific domains within those targets that give insights into disease mechanism and tractable therapeutic targets.

## Supporting information

Online methods section

## Acknowledgements

The Cambridge Human Research Tissue Bank is supported by the NIHR Cambridge Biomedical Research Centre. The sequencing was funded by the core grant from Wellcome Trust to the Wellcome Sanger Institute. MH is supported by a CRUK Programme Foundation Award (DRCPFA-Jun22\100001) and MRC Research Grant (MR/X00970X/1). SJM and JEC are supported by the MRC (MCMB MR/V028669/1, MCMB MR/Y011813/1) and the Alpha1 Foundation (pC ID: 830153).

---

**Supp. Figure 1.**
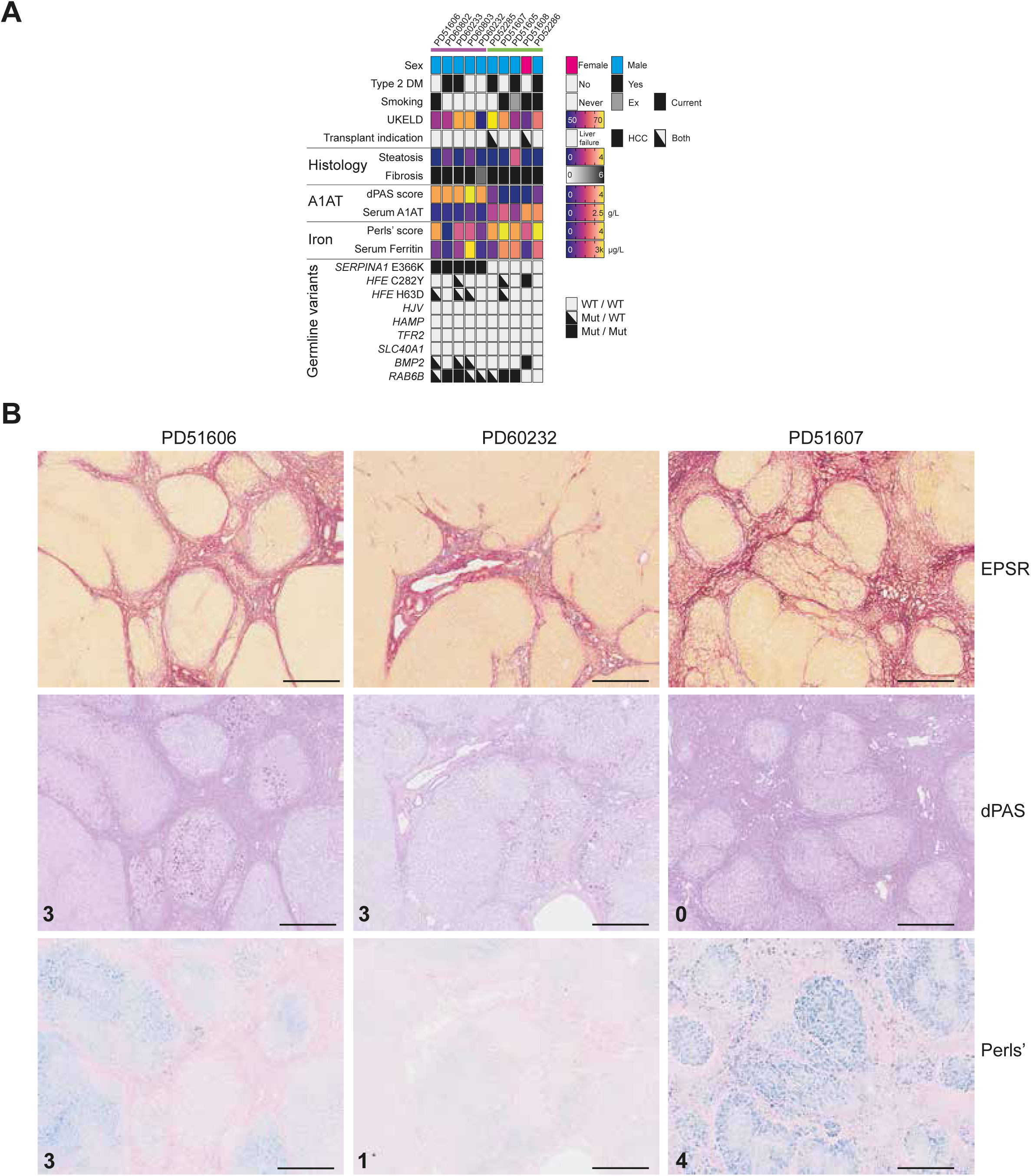
Clinical features and histopathology of patients included in the study. (A) Genomic and clinicopathological features of patients with alpha-1 anti-trypsin deficiency (A1AD, mauve header bar, n = 5) and haemochromatosis (green header bar, n = 5) included in the study. Germline variants were called from the exome or whole genome sequencing and reported for genes previously associated with either A1AD or haemochromatosis. (B) Example photomicrographs of serial liver sections stained for enhanced picosirius red (EPSR, upper panels), diastase-Periodic Acid Schiff (dPAS, middle panels) or Perls’ stain (lower panels) from three patients included in the study. Numbers on the panels are the dPAS granules / globules score (0-4) or hepatocellular iron deposition score (0-4) from global assessment of liver histology. Scale bars 1mm.

**Supp. Figure 2.**
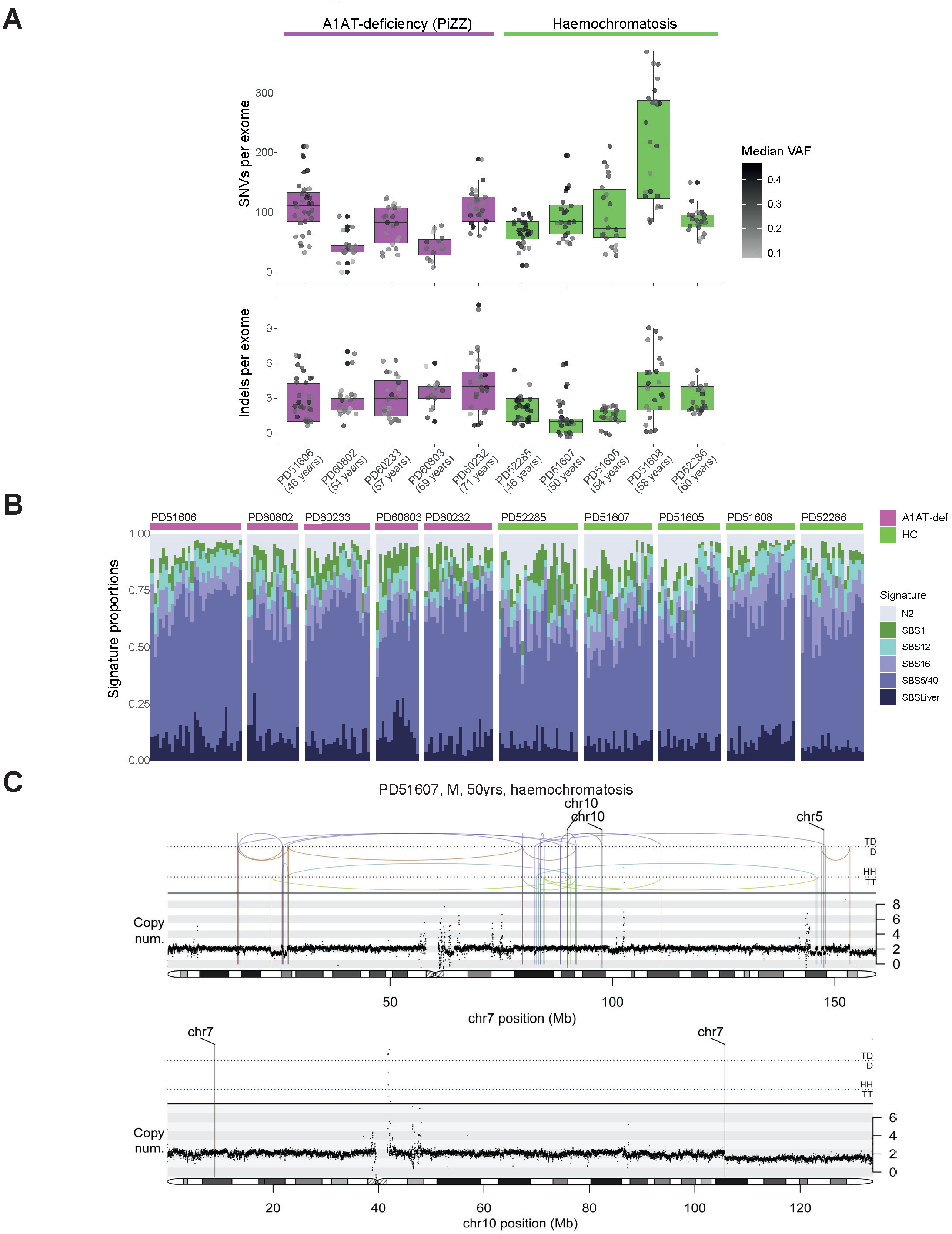
Mutational burden and large-scale rearrangements identified in the liver from A1AD and haemochromatosis. (A) Upper panel: burden of single-nucleotide variants (SNVs) per exome, corrected by sensitivity of mutation detection. Each box plot represents a patient (n = 10 patients; 305 microdissections) and each dot represents one laser-capture microdissected sample. The grey-to-black intensity of the points reflects the median variant allele fraction (VAF) of mutations in each microdissection. Boxes in the box plots indicate median and interquartile range; whiskers denote range. Lower panel: burden of indel variants per exome (n = 10 patients; 305 microdissections). (B) Relative proportional contributions of the six most abundant SBS signatures across A1AT-deficiency (purple-headed columns) and haemochromatosis (green-headed columns) donors. Each bar represents a laser capture microdissection, in ascending order of the number of coding mutations. (C) Chromothripsis involving chromosomes 5 and 10 (upper panel) and chromosomes 7 and 10 (lower panel), observed in patient with haemochromatosis (PD51607). Black points represent corrected read depth along the chromosome. Lines and arcs represent structural variants, coloured by the orientation of the joined ends (purple, tail-to-tail inverted; brown, head-to-head inverted; turquoise, tandem-duplication-type orientation; green, deletion-type orientation).

**Supp. Figure 3.**
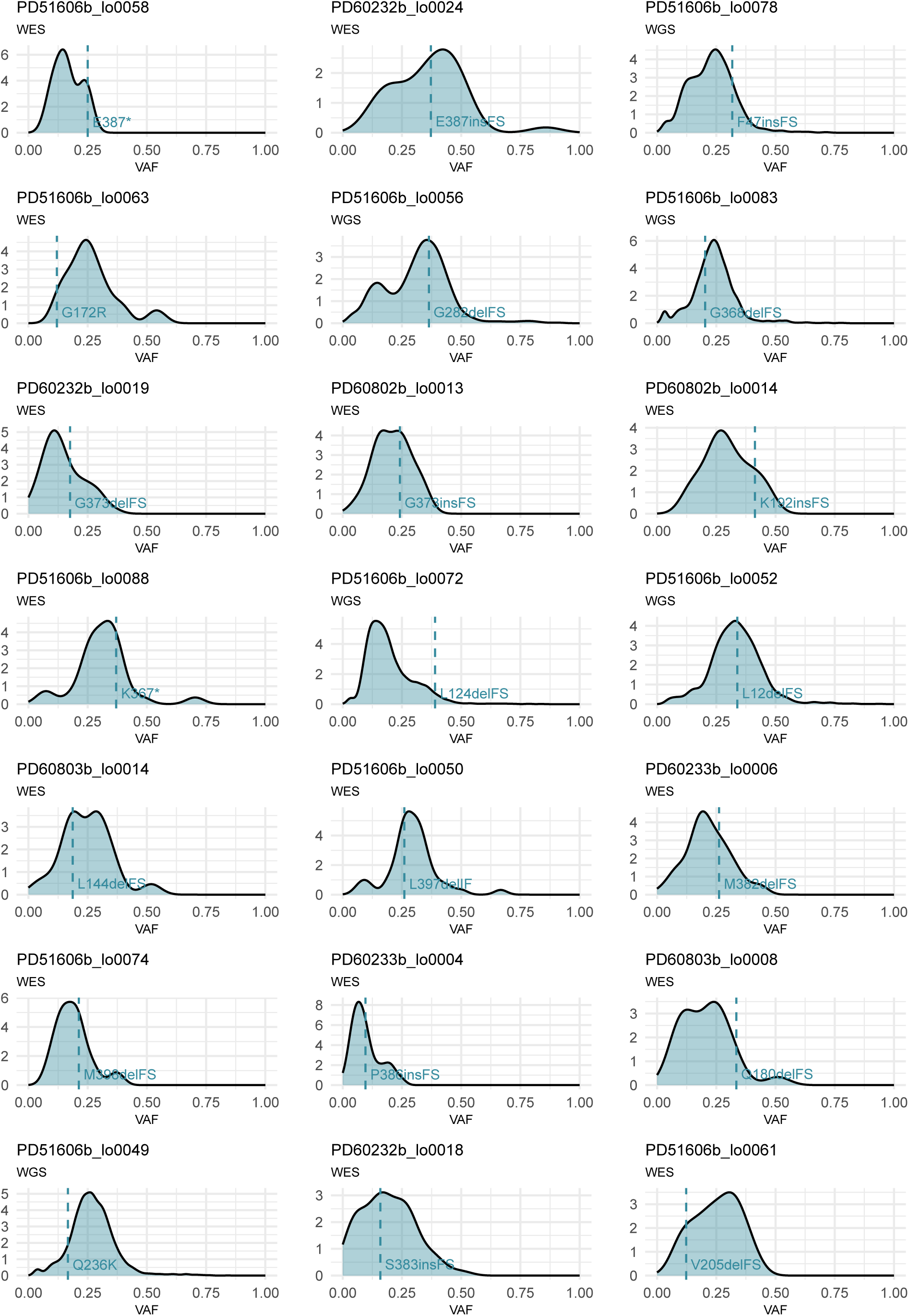
Variant allele fraction (VAF) density plots of LCM microdissections depicting VAFs of *SERPINA1* mutations. Dashed line drawn at the mean VAF of *SERPINA1* mutation in that sample. WES, whole exome sequencing. WGS, whole genome sequencing.

**Supp. Figure 4.**
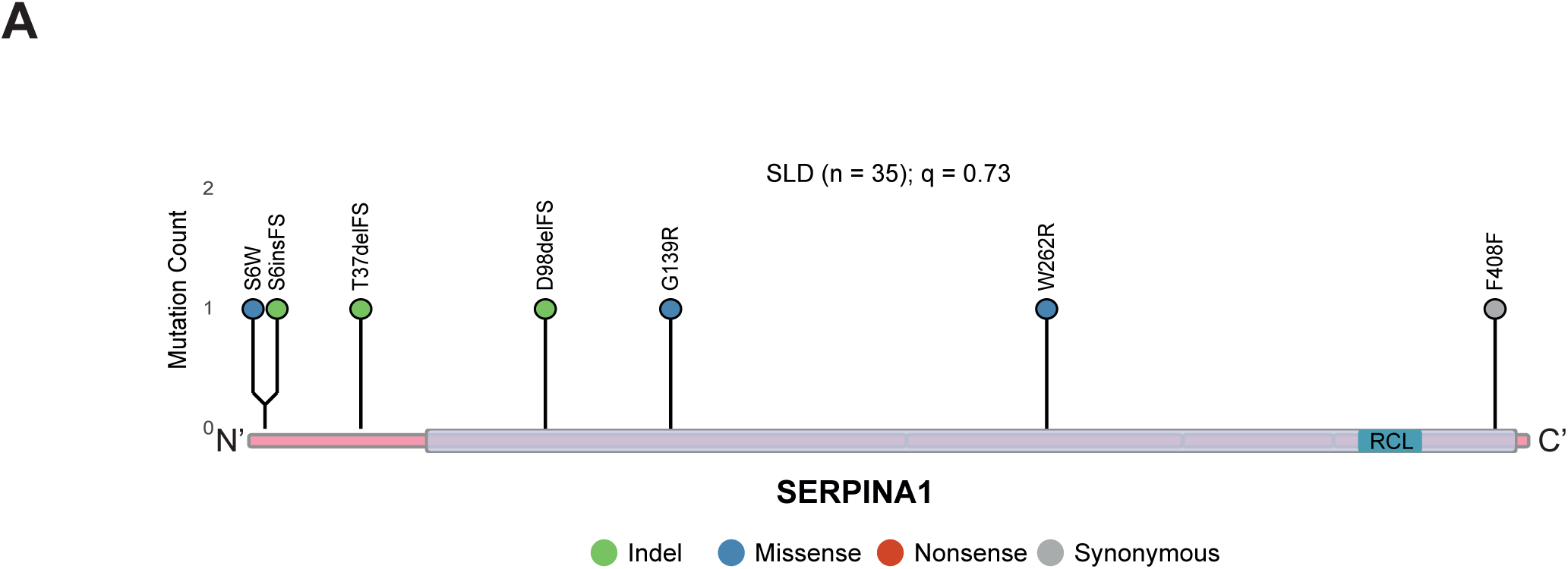
Somatic variants in *SERPINA1* and their protein coding consequences. (A) Distribution of variants detected in ***SERPINA1*** in liver samples from patients with SLD (n = 35) from Ng et al. Coding sequence of the gene represented on the x-axis, with exons in pink and protein domains in purple. Serpin, serine protease inhibitor domain; RCL, reactive centre loop.

**Supp. Figure 5.**
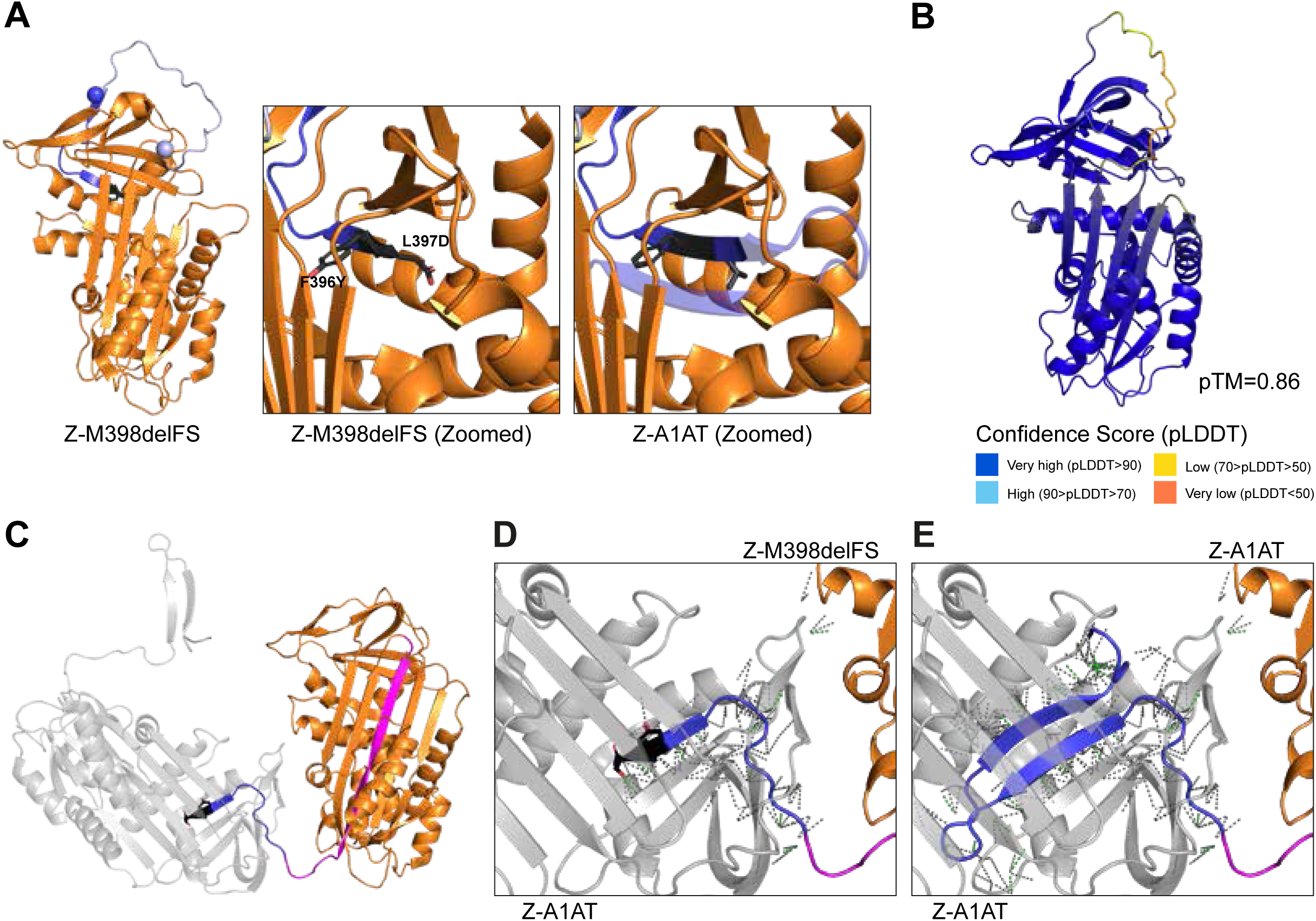
Structural prediction of the influence of Z-M398delFS somatic mutation. (A) Left: Cartoon of the primary sequence of the somatic A1AT mutant Z-M398delFS mapped as a portion of the structure of native full legth A1AT derived from Protein Data Bank (PDB):1QLP^17^. Magenta and blue regions reflect portions of the protein sequence lost in the somatic mutants K367* and E387* spespectively (mutation sites modeled as spheres), as shown in Figure 3C. Centre: Zoomed section of A1AT beta-sheet C showing the missense F396Y and L397D residues of Z-M398delFS in black with amino acid side chains rendered. Right: A comparative zoom of full length A1AT showing the side chains of native F396 and L397, with structure downstream of the Z-M398delFS premature STOP codon redendered opaque. (B) Alphafold3 prediction of the structure of A1AT Z-M398delFS with a predicted template modeling (pTM) score or 0.86, coloured by a per-atom confidence score (pLDDT). (C) Cartoon of the primary sequence of A1AT Z-M398delFS mapped onto the crystal structure of a Z-A1AT polymer from PDB:3T1P^18^, where Z-M398delFS is displayed as the donor protomer and the acceptor Z-protomer is grey. Residues downstream of K367 and E387 are shown in magenta and blue respectively. F396Y and L397D are shown in black. (D) Zoomed polymer interface shown from (C) showing contacts between protomers as dashed lines (polar contacts are green, other are grey). (E) A corresponding map of inter-protomer contacts between two molecules of full-length Z-A1AT.

**Supp. Figure 6.**
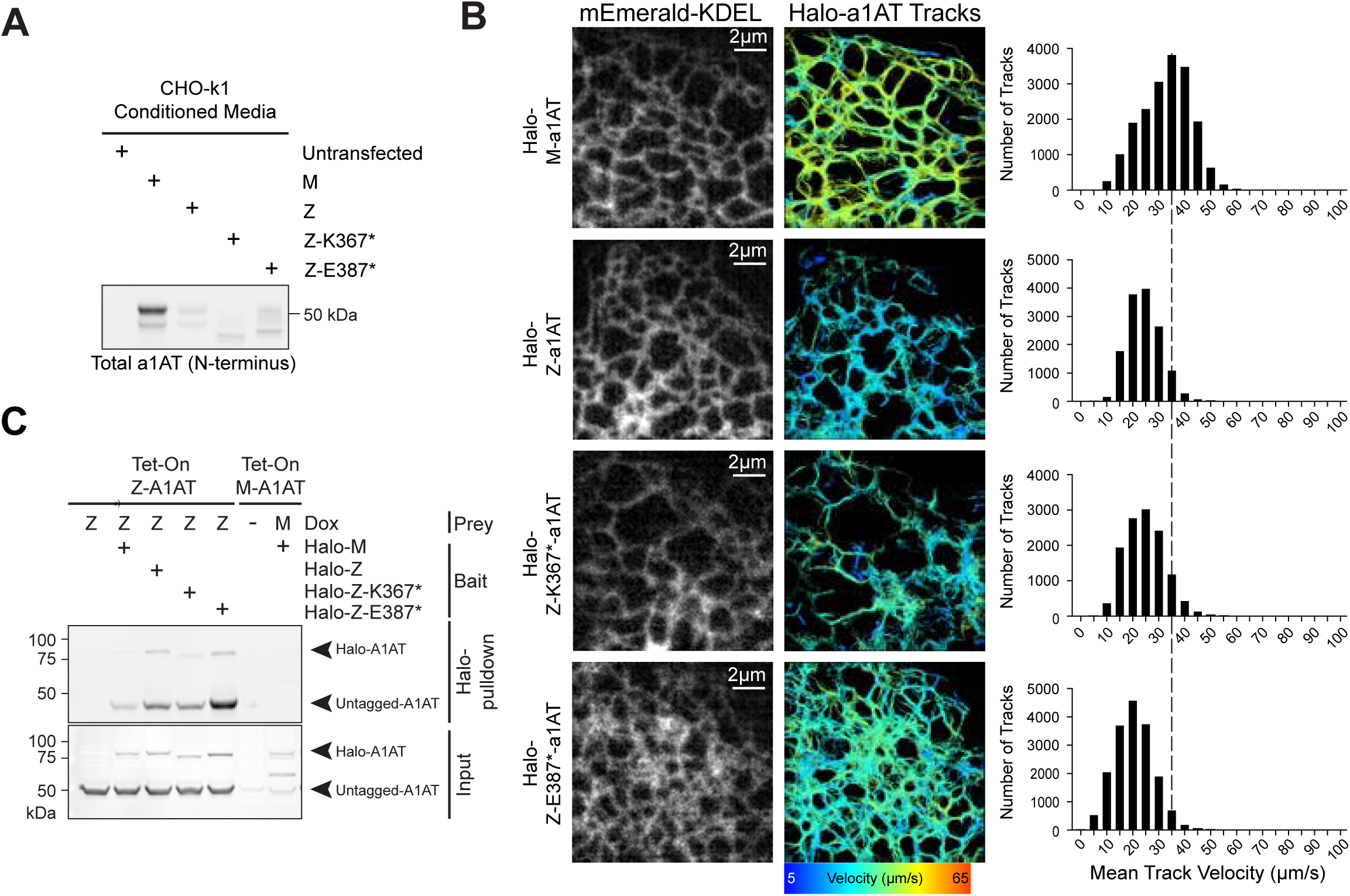
Somatic C-terminal truncation variants of *SERPINA1* do not rescue A1AT-secretion but have a short protein half-life. (A) Western blot analysis of A1AT secretion by immunoblotting of media conditioned for 24 hours by CHO cells transfected with the indicated A1AT-variant constructs. Immunoblotting was performed using a monoclonal antibody raised against an N-terminal fragment of human α1-antitrypsin (MA5-15521). (B) Micrographs of the ER marker protein mEmerald-KDEL expressed in COS7 cells (left images) with accompanying single-particle tracks of HaloTagged variants of a1AT labelled with PAJF646, colour coded by mean velocity (right images). Frequency distribution histograms show mean velocity of tracks analysed. A dashed line denotes the median velocity of M-a1AT. For each condition over 10,000 particle tracks were analysed across a minimum of 17 cells from 3 repeats. (C) CHO cells conditionally-expressing untagged Tetracycline-inducible (Tet-ON) Z-A1AT were transfected with expression vectors encoding HaloTagged variants of A1AT, 2 hours prior to 1µg/mL doxycycline treatment. Input and Halo-affinity purified samples were then separated by SDS-PAGE prior to immunoblotting with antibody recognising total (MA5-15521) A1AT.

**Supp. Figure 7.**
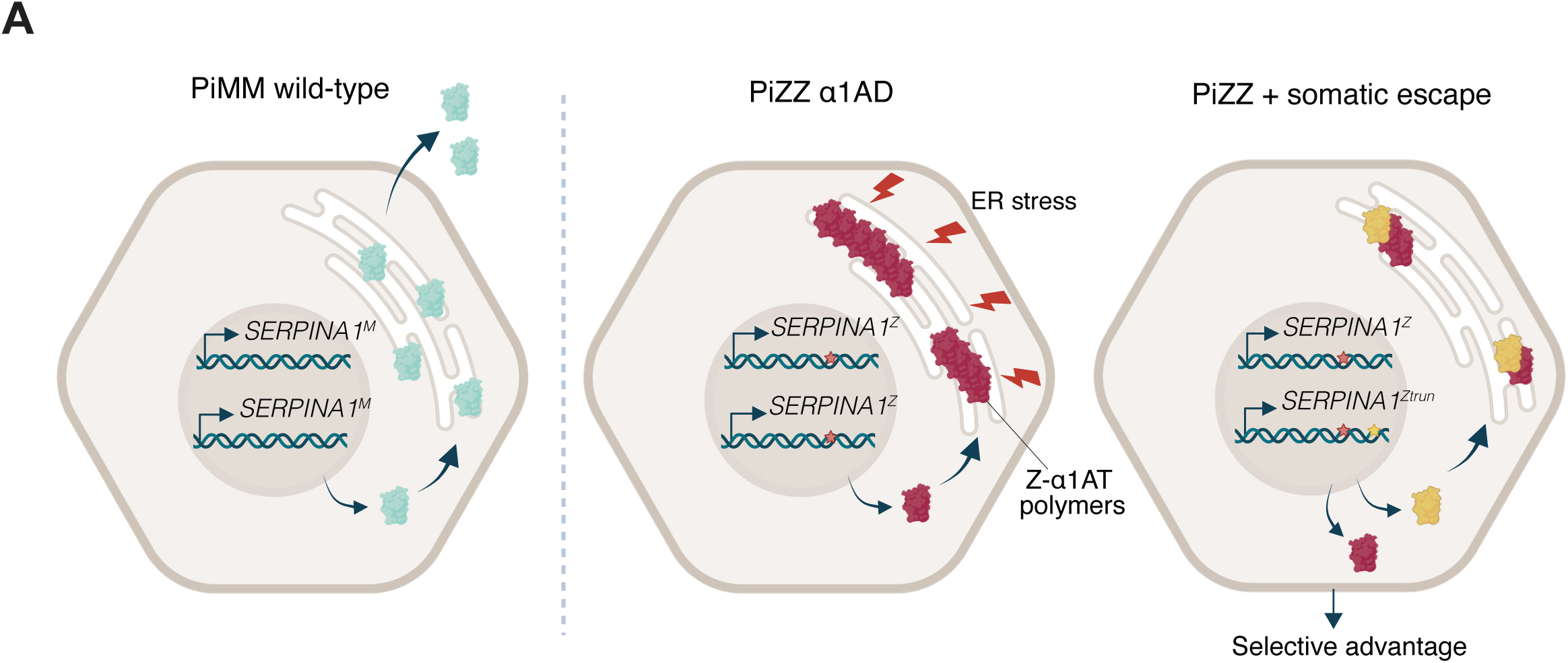
Model depicting cellular physiology of a1AT in differing genotypes. Homozygous MM-*SERPINA1* leads to expression, translation and secretion of functional A1AT protein; Homozygous ZZ-*SERPINA1* leads to translation and subsequent polymerisation of Z-A1AT, leading to ER disruption. Acquistion of a somatic escape variant in a single Z-*SERPINA1* allele leading to missense or truncation of the C-terminus reduces Z-A1AT polymerisation and ER disruption, providing a selective advantage compared to neighbouring hepatocytes with ZZ-*SERPINA1*.

## References

1. Moore, L. et al. The mutational landscape of human somatic and germline cells. Nature 597, 381–386 (2021).

2. Cagan, A. et al. Somatic mutation rates scale with lifespan across mammals. Nature 604, 517–524 (2022).

3. Zhu, M. et al. Somatic Mutations Increase Hepatic Clonal Fitness and Regeneration in Chronic Liver Disease. Cell 177, 608–621.e12 (2019).

4. Ng, S. W. K. et al. Convergent somatic mutations in metabolism genes in chronic liver disease. Nature 598, 473–478 (2021).

5. Brunner, S. F. et al. Somatic mutations and clonal dynamics in healthy and cirrhotic human liver. Nature 574, 538–542 (2019).

6. Huang, D. Q. et al. Global epidemiology of cirrhosis — aetiology, trends and predictions. Nat. Rev. Gastroenterol. Hepatol. 20, 388–398 (2023).

7. Berg, N. O. & Eriksson, S. Liver Disease in Adults with Alpha1-Antitrypsin Deficiency. N. Engl. J. Med. 287, 1264–1267 (1972).

8. Girelli, D. et al. Hemochromatosis classification: update and recommendations by the BIOIRON Society. Blood 139, 3018–3029 (2021).

9. Carrell, R. W. & Lomas, D. A. Alpha1-Antitrypsin Deficiency — A Model for Conformational Diseases. N. Engl. J. Med. 346, 45–53 (2002).

10. Wang, Z. et al. Positive selection of somatically mutated clones identifies adaptive pathways in metabolic liver disease. Cell (2023) doi:10.1016/j.cell.2023.03.014.

11. Jamialahmadi, O. et al. Exome-Wide Association Study on Alanine Aminotransferase Identifies Sequence Variants in the GPAM and APOE Associated With Fatty Liver Disease. Gastroenterology 160, 1634–1646.e7 (2021).

12. Ghouse, J. et al. Integrative common and rare variant analyses provide insights into the genetic architecture of liver cirrhosis. Nat. Genet. 1–11 (2024) doi:10.1038/s41588-024-01720-y.

13. Verweij, N. et al. Germline Mutations in CIDEB and Protection against Liver Disease. New Engl J Med 387, 332–344 (2022).

14. Martincorena, I. et al. Universal Patterns of Selection in Cancer and Somatic Tissues. Cell 171, 1029–1041.e21 (2017).

15. Ally, A. et al. Comprehensive and Integrative Genomic Characterization of Hepatocellular Carcinoma. Cell 169, 1327–1341.e23 (2017).

16. Faull, S. V. et al. The structural basis for Z α1-antitrypsin polymerization in the liver. Sci. Adv. 6, eabc1370 (2020).

17. Elliott, P. R., Pei, X. Y., Dafforn, T. R. & Lomas, D. A. Topography of a 2.0 Å structure of α1-antitrypsin reveals targets for rational drug design to prevent conformational disease. Protein Sci. 9, 1274–1281 (2000).

18. Yamasaki, M., Sendall, T. J., Pearce, M. C., Whisstock, J. C. & Huntington, J. A. Molecular basis of α1-antitrypsin deficiency revealed by the structure of a domain-swapped trimer. EMBO Rep. 12, 1011–1017 (2011).

19. Dickens, J. A. et al. The endoplasmic reticulum remains functionally connected by vesicular transport after its fragmentation in cells expressing Z-α1-antitrypsin. FASEB J. 30, 4083–4097 (2016).

20. Chambers, J. E. et al. Z-α1-antitrypsin polymers impose molecular filtration in the endoplasmic reticulum after undergoing phase transition to a solid state. Sci. Adv. 8, eabm2094 (2022).

21. Segeritz, C.-P. et al. hiPSC hepatocyte model demonstrates the role of unfolded protein response and inflammatory networks in α1-antitrypsin deficiency. J. Hepatol. 69, 851–860 (2018).

22. Ordóñez, A. et al. Endoplasmic reticulum polymers impair luminal protein mobility and sensitize to cellular stress in alpha1-antitrypsin deficiency. Hepatology 57, 2049–2060 (2013).

23. Grimm, J. B. et al. A general method to improve fluorophores for live-cell and single-molecule microscopy. Nat. Methods 12, 244–250 (2015).

24. Tinevez, J.-Y. et al. TrackMate: An open and extensible platform for single-particle tracking. Methods 115, 80–90 (2017).

25. Ronzoni, R. et al. The molecular species responsible for α1-antitrypsin deficiency are suppressed by a small molecule chaperone. FEBS J. 288, 2222–2237 (2021).

26. Lieberman, J., Mittman, C. & Gordon, H. W. Alpha1 Antitrypsin in the Livers of Patients with Emphysema. Science 175, 63–65 (1972).

27. Behrens, M. A. et al. The Shapes of Z-α 1-Antitrypsin Polymers in Solution Support the C-Terminal Domain-Swap Mechanism of Polymerization. Biophys. J. 107, 1905–1912 (2014).

28. Wooddell, C. I. et al. Development of an RNAi therapeutic for alpha-1-antitrypsin liver disease. JCI Insight 5, (2020).

29. Strnad, P. et al. Fazirsiran for Liver Disease Associated with Alpha1-Antitrypsin Deficiency. N. Engl. J. Med. 387, 514–524 (2022).

30. Robinson, P. S. et al. Increased somatic mutation burdens in normal human cells due to defective DNA polymerases. Nat Genet 53, 1434–1442 (2021).

31. Lee, B. C. H. et al. Mutational landscape of normal epithelial cells in Lynch Syndrome patients. Nat. Commun. 13, 2710 (2022).

32. Hirschhorn, R. et al. Spontaneous in vivo reversion to normal of an inherited mutation in a patient with adenosine deaminase deficiency. Nat. Genet. 13, 290–295 (1996).

33. Burrow, K. L. et al. Dystrophin expression and somatic reversion in prednisone-treated and untreated Duchenne dystrophy. Neurology 41, 661–666 (1991).

34. Klein, C. J. et al. Somatic reversion/suppression in Duchenne muscular dystrophy (DMD): evidence supporting a frame-restoring mechanism in rare dystrophin-positive fibers. Am. J. Hum. Genet. 50, 950–9 (1992).

35. Demers, S. I., Russo, P., Lettre, F. & Tanguay, R. M. Frequent mutation reversion inversely correlates with clinical severity in a genetic liver disease, hereditary tyrosinemia. Hum. Pathol. 34, 1313–1320 (2003).

36. Tan, S. et al. Somatic genetic rescue of a germline ribosome assembly defect. Nat. Commun. 12, 5044 (2021).

