## Supplementary material for "Convergent evolution of somatic escape variants in *SERPINA1* in the liver in alpha-1 anti-trypsin deficiency": Online methods section

### Materials and methods

#### Liver samples

All liver samples were collected with written informed consent from Addenbrooke's Hospital, Cambridge, UK, according to procedures approved by Local Research Ethics Committees (20/NI/0109, 16/NI/0196). All participants consented to publication of research results. The liver samples were snap-frozen in liquid nitrogen and stored at −80 °C in the Human Research Tissue Bank of the Cambridge University Hospitals NHS Foundation Trust.

Clinical details, anthropometrics and blood results are detailed in Supplementary Fig 1 and Supplementary Table 1. Background diseased liver tissue was obtained from individuals with A1AT deficiency or haemochromatosis, undergoing liver transplantation for HCC or liver failure (Supplementary Table 1). Patients were identified from clinical history, pre-operative investigations and explant histology. All patients with haemochromatosis had current or previously elevated serum ferritin with significant hepatocellular siderosis by Perls' stain. Pre-operative clinical genotyping was only performed for the common C282Y and H63D variants of the *HFE* gene. A1AT deficiency patients were diagnosed based on PiZZ A1AT electrophoretic phenotype and sub-normal serum A1AT levels by routine clinical testing. The *SERPINA1, HFE, HJV, HAMP, TFR2, SLC40A1, BMP2, RAB6B* genotypes[1] were derived from the whole-genome (WGS) or whole-exome sequencing data (Supplementary Fig 1 and Supplementary table 2). Clinical details and sequencing data from patients with steatotic liver disease was derived from previous publications [2,3].

The explant liver histology was reviewed by a specialist liver histopathologist (A.D., blinded to the other results in the study). Histological features from the liver specimens were scored according to the Kleiner[4] system on formalin-fixed paraffin-embedded (FFPE) samples away from the fresh-frozen block used for the laser-capture microdissection (LCM). The Kleiner score, developed for NAFLD, assesses the presence of steatosis, lobular inflammation and hepatocyte ballooning to generate a cumulative NAFLD activity score (NAS); artefactual inflammation secondary to surgical handling was excluded. We applied this to the diseased samples, which, in the absence of a validated scoring system for A1AT deficiency or haemochromatosis, allows comparability between all study samples regardless of disease aetiology. The histological findings were scored for siderosis by Perls' staining using the Scheuer system[5] and dPAS-positive globule deposition (Supplementary Fig 1) as previously described[6]. Fibrosis was assessed using both the Kleiner[4] and the Ishak[7] scoring systems. The presence or absence of cellular or nodular dysplasia was assessed globally in clinical FFPE samples (Supplementary Table 1).

**Sample preparation**

The protocols for preparing liver tissue sections, LCM and subsequent cell lysis, DNA extraction, and WGS were previously described[3,8]. In brief, for 6 biopsies (PD51605b, PD51606b, PD51607b, PD51608b, PD52285b, PD52286b), 20-μm-thick tissue sections (prepared with a Leica cryotome) were fixed with 70% ethanol. The other 4 biopsies (PD60232b, PD60233b, PD60802b, PD60803b) were fixed in PAXgene solutions (PreAnalytiX), processed using a Tissue Tek VIP 6 AI tissue processor (Sakura Finetek, Netherlands), embedded in paraffin and 16-μm-thick sections generated using an Accu-Cut SRM 200 microtome (Sakura Finetek). All LCM sections were mounted on polyethylene naphthalene (PEN) membrane glass slides (Leica Microsystems) and stained with H&E for subsequent LCM generation using a Leica Microsystems LMD 7000).

**Laser-capture microdissection**

For 6 biopsies (PD51605b, PD51606b, PD51607b, PD51608b, PD52285b, PD52286b), 48 microdissections were cut with a target area of 86,000 $\mu m^2$, with the same x,y-region cut into the same well from two adjacent z-stacks. For the remaining 4 biopsies (PD60232b, PD60233b, PD60802b, PD60803b), cuts were taken at 166,000 $\mu m^2$ without z-stacking. Overall, microdissection sizes corresponded to 800-1000 hepatocytes. Images were taken before and after LCM. The micro-dissected samples were then lysed using the Arcturus PicoPure DNA Extraction Kit (Thermo Fisher Scientific) following the manufacturer's instructions. DNA libraries for Illumina sequencing were prepared using a protocol optimized for low input amounts of DNA [9]. Resulting libraries were submitted for paired-end WGS or exome sequencing.

**Exome sequencing**

Exome capture was performed using either SureSelect All Exon v5 bait set (Agilent, S04380110) or Twist Human Core Exome (Twist Biosciences) bait set. Samples were multiplexed and sequenced using 150 bp paired-end reads on an Illumina NovaSeq 6000 (average pool size of 40). Paired-end reads were aligned to human genome assembly GRCh38 using BWA-MEM[10]. Duplicate reads were marked using biobambam[11] and sample contamination estimates were calculated using VerifyBamID[12]. Library complexity and coverage statistics were calculated using Picard (http://broadinstitute.github.io/picard/). The median on-target coverage across all samples and genes was 41´. Across donors, median coverage ranged from 28´ (PD60802b) to 48´ (PD51608b).

**Whole-genome sequencing**

Whole-genome sequencing (WGS) was performed using 150 bp paired-end reads on an Illumina NovaSeq 6000. DNA sequences were aligned to the GRCh38 reference genome using BWA-MEM. Duplicate reads were marked using biobambam[11]. The calculation of library statistics was performed using the CollectWgsMetrics function of Picard Tools, rather than the CollectHsMetrics function used for exome sequencing data.

**Calling of SNVs and indels from exome and WGS data**

SNVs were called using the CaVEMan (cancer variants through expectation maximisation) algorithm[13] against an *in silico* unmatched normal: a BAM file generated from the human reference genome (GRCh38). Indel calling was performed using cgpPindel[14]), with filtering strategies the same as for SNVs. Duplicate reads and LCM library preparation-specific artefactual variants resulting from the incorrect processing of secondary cruciform DNA structures were removed with bespoke post-processing filtering[9]. Only variants with at least two supporting sequencing reads were considered. To filter shared artefacts, we applied a beta-binomial-based filtering approach as previously described[15]. Putative germline variants were filtered using a one-sided exact binomial test used on the aggregated counts of reads supporting the variant and the total depth at that site, as described previously[15]. This tests whether the observed variant counts are likely to have come from a germline distribution or from a distribution with a lower true VAF (probably somatic). For sex chromosomes in male individuals, the binomial probability (true VAF) for comparison was set to 0.95 rather than 0.5. The resulting P values are corrected for multiple testing using a Benjamini–Hochberg correction. Any variant with a q value < $10^{-5}$ was categorized as a putative somatic variant.

#### Structural variant calling from WGS data

Structural variants (SVs) were called with GRIDDS[16] (version 2.9.4), used with default settings. SVs larger than 1kb in size with QUAL ≥ 250 were included. For SVs smaller than 30kb, only SVs with QUAL ≥ 300 were included. Furthermore, SVs that had assemblies from both sides of the breakpoint were only considered if they were supported by at least four discordant and two split reads. SVs with imprecise breakends (i.e., the distance between the start and end positions > 10bp) were filtered out. We further filtered out SVs for which the standard deviation of the alignment positions at either ends of the discordant read pairs was smaller than five. To remove potential germline SVs and artefacts, we generated the panel of normal by adding in-house normal samples (n = 350) to the GRIDSS panel of normal. SVs found in at least three different samples in the panel of normal or in matched normals were removed.

#### Copy number variant calling from WGS data

Somatic copy-number variants (CNVs) were called using the Allele-Specific Copy number Analysis of Tumours (ASCAT) algorithm[17] as part of the ascatNGS package[18] (https://github.com/ Crick-CancerGenomics/ascat). ASCAT was run with default parameters with the exception of a segmentation penalty of 100. A bespoke filtering algorithm -

ascatPCA - was used to reduce the number of false-positive calls that can arise when analysing genome sequences from normal tissue (https://github.com/hj6-sanger/ascatPCA). ascatPCA extracts a noise profile by aggregating the LogR ratio from across a panel of normal unrelated samples and subtracts this signature from that observed in the sample being analysed using principal component analysis.

##### Extraction of mutational signatures

As previously[2,3], The HDP algorithm as implemented in the HDP R package (https://github.com/nicolaroberts/hdp), was used to extract mutational signatures, based on a reference catalogue of 65 previously identified 192-context-based mutational signatures from the Pan-Cancer Analysis of Whole Genomes (PCAWG) study[19].

##### Bayesian Dirichlet process for clustering VAFs across multiple samples

We used the nonparametric Bayesian hierarchical Dirichlet process (HDP) to group SNVs based on similar variant allele fractions (VAFs) identified across multiple microdissections in each patient biopsy. This method, called N-dimensional Dirichlet Process (NDP) clustering, has been detailed previously[3]. We ran the algorithm with 15,000 burn-in iterations, followed by 25,000 iterations of posterior Gibbs sampling for the clustering process. During each iteration, there is a defined probability that mutations will be allocated to new clusters that did not exist in the previous iteration. Existing clusters can also be eliminated if member mutations are reassigned to another cluster. This adaptive process allows for the dynamic adjustment of the number of clusters throughout the sampling. To prevent the formation of uninformative clusters, we capped the number of SNV clusters at 100 per patient. We also employed a multi-threaded version of the ECR algorithm, adapted from the label.switching R package (DOI: 10.18637/jss.v069.c01), to correct for label switching efficiently. For subsequent analyses, we only considered SNV clusters that included at least 50 distinct mutations.

#### Construction of phylogenetic trees

In this study, we applied the statistical pigeonhole principle[20] to deduce the phylogenetic clonal relationships among SNV clusters identified in each patient by the NDP algorithm. Here, each cluster forms a branch on a phylogenetic tree. Evidence that a cluster is considered nested within another is considered strong if its mutation-carrying cell fraction (CF) is consistently lower than that of another cluster across all sampled microdissections, and if the combined mutant CFs exceed 100%. A combined CF of ≤ 100% indicates only weak evidence of such nesting. If only certain microdissections show a lower CF for one SNV cluster compared to another, these clusters are deemed independent, not nested within one another. Our analysis was restricted to clusters with a mutant CF greater than 0.05. We calculated the CF for each SNV cluster by doubling the median VAF in each microdissection, assuming diploidy. SNV clusters with microdissections lacking shared mutations with others in the same cluster were divided into new, independent clusters. These were then reassessed for their phylogenetic relationships to all other clusters from the same patient biopsy, using the pigeonhole principle. Additionally, a naive Bayes algorithm was employed to categorize each identified indel into the SNV clusters detected by the NDP algorithm.

#### Analysis of driver variants

To determine whether any coding variants were under selection in diseased liver tissues, the dN/dScv method[21] on the gene level was used. The algorithm identifies genes with an excess of non-synonymous mutations relative to the expected number from the synonymous mutation rate. For this analysis, variants called from whole genomes were collapsed to unique events per SNV cluster identified by the NDP algorithm. Any variants identified in exome data which were not already called in whole genomes from the same patient were collapsed to unique mutations per individual and added to the dN/dScv input. Mutations with q values of < 0.1 were considered to be under positive selection.

**Extraction of mutational signatures from SNV contexts using HDP**

To identify possibly undiscovered mutational signatures in the liver from α1-antitrypsin deficiency and haemochromatosis patients, the hierarchical Dirichlet process (HDP, see https://github.com/nicolaroberts/hdp) was ran on the 96 trinucleotide counts of all microdissected samples, collapsed to unique mutations across samples. To avoid over-fitting, samples with fewer than 50 mutations were not included in the signature extraction. HDP was run with individual patients as the hierarchy, in twenty independent chains, with a burn-in of 20,000 and the collection of 100 posterior samples off each chain with 200 iterations between each. Due to the lack of novel signatures in this data set, the remainder of mutational signature analysis was performed by fitting the identified set of signatures from HDP to trinucleotide counts from each microdissection using the R package sigfit (DOI: https://doi.org/10.1101/372896).

**Protein structure prediction modelling**

Structural predictions of the Z-K367*, Z-E387*, and Z-M398delFS somatic mutant A1AT proteins were performed using Pymol molecular visualisation software (Version 2.5.2, Schrodinger) to map amino acid sequences onto the crystal structure of native human A1AT (pdb:1QLP)[22], identifying secondary structures lost in somatic mutant proteins. To predict the ability of somatic mutant proteins to form mixed polymers with full length Z-A1AT, their sequences were mapped onto a single protomer within the crystal structure of a human Z-A1AT polymer (3T1P)[23]. Atoms of the 3T1P protomer that are lost through truncation of somatic mutants were removed from the displayed image, to highlight any incompatibility with polymerisation. The two missense residues in Z-M398delFS (F396Y, L397D) were modelled using the Pymol mutagenesis function, selecting side chain rotamers of Y396 and D397 that best conserved the resolved density in 3T1P. Somatic mutant structural models did not undergo energy minimisation or geometry optimisation processes, and hence are used only to predict feasibility of polymerisation and not changes to the protein fold. Structural prediction of Z-M398delFS A1AT was carried out using Alphafold 3[24], where the top-ranked model is displayed as a cartoon coloured by pLDDT score.

**Cloning and constructs**

Somatic variants of Z-$\alpha_1$-antitrypsin (Z-A1AT) (Z-K367*, Z-E387*) were generated by site directed mutagenesis of a Z-A1AT coding sequence (untagged and N-terminally tagged with HaloTag) used previously[25]. Bicistronic expression vectors were generated from pGL4.2 encoding A1AT tagged with Halo-Tag (M-, Z-, Z-K367*-, Z-E387*-A1AT variants) mutant and the ER marker protein moxsynGFP-KDEL by Gibson assembly (New England Biolabs). Sequencing of the plasmids was carried out at Plasmidsaurus (Oxford Nanopore Technologies).

#### Sequences of primers used in the study

| | |
|---|---|
| pGL4.2_ER-mox_1362-761_fwd | CCAAAAATAATGATCTAGAACCGGTCATGG |
| pGL4.2_ER-moxGFP_1362-761_rev | ATAGCAGCATGGTGGCGCTAGTGTCAGAAG |
| pHalo-Z_597-2834_fwd | TAGCGCCACCATGCTGCTATCCGTGCCG |
| pHalo-Z_597-2834_rev | TTCTAGATCATTATTTTTGGGTGGGATTCACCAC |
| A1AT_K367X_fwd | GCAGCTTCAGTCCCTTACTTGTCGATGGTCAGC |
| A1AT_K367X_rev | GCTGACCATCGACAAGTAAGGGACTGAAGCTGC |
| A1AT_E387X_fwd | TTTGTTGAACTTGACCTAGGGGGGGATAGACATGG |
| A1AT_E387X_rev | CCATGTCTATCCCCCCCTAGGTCAAGTTCAACAAA |

#### Mammalian cell culture

Chinese Hamster Ovary (CHO) parental cells were cultured in F12 Ham's nutrient mixture (Merck, Germany) supplemented with 10% fetal bovine serum (FBS) and 2mM GlutaMAX (Thermo Fisher Scientific, USA) at 37°C, 5% $CO_2$. CHO Tet-ON cells with inducible expression of Z-A1AT were cultured in high glucose DMEM (Merck, Germany) supplemented with 10% tetracycline-free fetal bovine serum (FBS), 2mM GlutaMAX, and non-essential amino acids (Thermo Fisher Scientific, USA). COS7 cells (MERCK, Germany) were cultured in high glucose DMEM supplemented with 10% fetal bovine serum (FBS) and 2mM GlutaMAX.

#### Live-cell imaging

CHO parental cells were seeded at a density of 8.1 x $10^4$ cells/$cm^2$ in 6 cm dishes and transfected the same day with a bicistronic vector encoding moxGFP-KDEL and HaloTag-A1AT (M, Z, Z-K367*, Z-E387* variants). Transfection was performed according to the Lipofectamine LTX reagent protocol (Life Technology, UK). For one

6 cm dish transfection, 560 µl of Opti-MEM were mixed with 1.5 µg of DNA. Lipofectamine was then added dropwise to the surface of the solution in a ratio of 4ul per µg of DNA (6 µl per one dish). The resulting solution was then mixed by shaking and incubated at room temperature for 15 minutes prior to addition to cells. 24 hours later, cells were trypsinised and sorted for a GFP-positive population using the BD Influx™ Cell Sorter, gating around the median fluorescence intensity. After sorting, cells were seeded at $1x10^4$ cells/$cm^2$ in an 8-well ibidi slide. Imaging was performed 48 h after seeding. Prior imaging cells were labelled with 0.5µm JFX549 (Janelia Fluor) in Opti-MEM for 15 min at 37°C. Airyscan images were collected using a 63x oil objective with a numerical aperture of 1.4, on an LSM 880 (Zeiss, Germany) confocal microscope using Airyscan detector and processed using the Zen 2.6 software package (Black edition, Zeiss, Germany. Quantitation of ER morphology was performed using standard confocal imaging to avoid unnecessarily large file sizes. Cells were imaged using 488 nm and 561 nm lasers. ImageJ (Fiji) software package was used to process images, applying a gamma correction factor of 0.65 to Airyscan images in order to simultaneously visualise ER inclusions and tubules. Cells were counted based on ER morphology, categorised as either reticular ER or as containing ER-inclusions. Quantitation was performed using the moxGFP-KDEL channel, to avoid bias introduced by variable accumulation (and hence fluorescence intensity) of different A1AT variants.

**Single particle tracking**

Single particle tracking of HaloTag-A1AT variants was performed as described previously[26]. Briefly, COS7 cells were seeded at $4.16x10^3$ cells/$cm^2$ on Matrigel coated No. 1.5H glass coverslips (Marienfeld, Germany). Cells were transfected 4 hours later using 1ug DNA and 3ul Fugene 6 (Promega, US) as per manufacturer's instructions, to drive expression of an ER marker protein (mEmerald-KDEL) and HaloTag-A1AT variants. Cells were imaged 18 hours after transfection, following labelling with PA-JF646 photoactivatable ligand[27]. Images were acquired using a Zeiss Elyra7 wide-field microscope using a ×63 1.46 NA oil immersion TIRF objective in HiLo modality. Synchronous capture of two fluorescence channels was performed using an OptoSplit beam splitter (Cairn Research Ltd., UK) and two pco.edge sCMOS cameras (PCO, Germany). Excitation of fluorophores was carried out at 488 nm (mEmerald) and 642 nm (HaloTag PA-JF646). Images were captured with an exposure time of 4ms at a frame rate of 167 Hz. Photoactivation of PA-JF646 bound to HaloTag-KDEL was achieve by continuous exposure to 405-nm laser light to tune an appropriate density of particles within the 128 × 128 pixel frame to permit tracking. A Laplacian of Gaussian filter was applied to HaloTag-A1AT particle images, and single spot detection was carried out with subpixel localization. A simple Linear Assignment Problem (LAP) tracking function of the TrackMate ImageJ plugin[28] was applied to particle image series, with a threshold minimum number of spots per track set to 30. Mean track velocity was extracted for each assigned particle track and collated with equal weighting across all cells in an experimental group.

**Sandwich enzyme-linked immunosorbent assay (ELISA) for α1-antitrypsin (A1AT)**

Parental CHO cells were seeded at a density of 2.08x$10^6$ cells/$cm^2$ in 6-well cell culture plates (Greiner Bio-One, UK). Transfection was performed as described above for CHO parental cells. Cell culture media was substituted with 1mL of Opti-MEM reduced serum medium (ThermoFisher Scientific, USA) 24 hours after transfection.

Conditioned Opti-MEM and cell lysates were collected 48 hours after transfection. Cells were harvested by washing twice with PBS on ice. 200mL of ice-cold lysis buffer (0.01M 4-(2-hydroxyethyl)-1-piperazine ethane sulfonic acid (HEPES), 0.05M NaCl, 0.56M sucrose, 0.1mM EDTA, 0.5% v/v Triton) was applied to each well, cell homogenates were collected and centrifuged at 16,100xg for 10 minutes at 4°C in a bench-top centrifuge. The lysate supernatant was used for the subsequent steps of ELISA. ELISA plates (96-well, COSTAR Corning) were coated overnight at room temperature in a humidity chamber with a rabbit polyclonal antibody (ab9373, Abcam) raised against A1AT in PBS. The plates were washed three times with washing buffer (0.9% w/v NaCl, 0.05% v/v Tween20) and subsequently blocked in blocking buffer (PBS with 0.25% w/v BSA, 0.05 %v/v Tween20, 0.1 % sodium azide) for 1 hour at room temperature. To generate standard curves, Z-A1AT monomer of known concentrations was used in a dilution series. Plates were incubated for 2 hours at room temperature with test samples. Plates were subsequently washed 3 times and all liquid removed before addition of 50μL of primary monoclonal antibody $mAb_{3C11}$ (HycultBiotech, USA) in blocking buffer to each well, and incubated for 2 hours at room temperature. Plates were washed and incubated with rabbit anti-mouse horseradish peroxidase (HRP)-labelled antibody (a9044, Sigma-Aldrich), diluted 1:20,000 in blocking buffer without sodium azide. 5μL of the HRP substrate solution (Sigma-Aldrich, UK) was applied and plates were developed in the dark at room temperature for 10 minutes. The reaction was stopped by adding 50μL of 1M $H_2SO_4$. Absorbance was measured at 450nm wavelength in a Tecan Speak plate reader. Data analysis was performed in Microsoft Excel. Standard curves were plotted from absorbance values of known concentrations of Z-A1AT. The linear part of the standard curves was used to determine concentration of A1AT in analysed test samples.

**Native- and SDS- polyacrylamide gel electrophoresis (PAGE)**

Parental CHO and Tet-On CHO cells were seeded at a density of $2.08x10^6$ cells/$cm^2$, transfected as described above for CHO parental cells, and grown in 6-well cell culture plates (Greiner Bio-One, UK) to approximately 90% confluency for a further 48 hours. Cells were washed twice with PBS on ice. 100 µL of ice-cold lysis buffer were applied to each well, and cell homogenates were collected. Cell lysates were sonicated in a water bath sonicator for 30 minutes at 4°C, centrifuged at 16,100xg for 10 minutes at 4°C in a bench-top centrifuge and supernatant (soluble lysate) was separated from pelleted material before snap freezing. Pelleted material was resuspended in 100µL of lysis buffer, sonicated in a water bath sonicator for 45 minutes at 4°C (insoluble lysate). For Native PAGE, sonicated soluble lysate containing 80µg of protein, or an equivalent volume of the corresponding insoluble lysate, was mixed with 6 µL of native loading buffer (50% v/v glycerol, 0.01% bromophenol blue) and loaded onto an acrylamide native gel (resolving gel composition: 7.5% w/v Acrylamide-bisacrylamide mixture (37.5:1), 0.37mM Tris-HCl pH8.8, 0.12% w/v APS, 0.2% v/v TEMED. Stacking gel composition 5.3% w/v acrylamide-bisacrylamide mixture (37.5:1), 110 mM Tris-HCl pH6.8, 0.125% w/v APS, 0.15% v/v TEMED). Under native conditions, samples were run at 100V until the dye front reached the bottom of the gel, then run for an additional 30 minutes. Anode buffer (0.1M Tris-HCl pH7.8) and cathode buffer (50mM Tris-HCl pH8.9, 68mM glycine) were used. For SDS-PAGE, soluble lysate containing 80µg of protein, or an equivalent volume of the corresponding insoluble lysate, were prepared in SDS-loading buffer (312.5mM Tris-HCl pH6.8, 50% v/v glycerol, 10% w/v SDS, 0.05% w/v bromophenol blue, 50mM DTT) and were heated at 75°C for 10 minutes before loading on a 10% acrylamide gel. Native-PAGE gels were transferred onto nitrocellulose membranes using a wet transfer system with Native transfer buffer (20 mM Tris and 140 mM glycine) for 2 hours at 200mA. SDS-PAGE gels were transferred in transfer buffer (25mM Tris-HCl pH8.3, 250mM glycine, 20% v/v methanol) for 1.5 hours at 300mA. Membranes were blocked in 5% milk dissolved in 1x Tris buffered saline (TBS) solution and incubated overnight at 4°C in the primary antibody. Membranes were then incubated with the appropriate secondary antibody (Li-Cor Biotechnology, US) at room temperature (RT) for 2 hours (Native-PAGE), or 1 hour (SDS-PAGE) prior to detection blot detection using a Li-Cor CLx scanner. A monoclonal $\alpha_1$-antitrypsin antibody (MA5-15521, Thermo Fisher Scientific) raised against a peptide consisting of amino acids 40-184 of the human protein was used for SDS-PAGE western blotting. A polyclonal antibody raised against full-length human $\alpha_1$-antitrypsin (A0409, Sigma-Aldrich) was used for Native-PAGE western blot (total $\alpha_1$-antitrypsin pool). The human $\alpha_1$-antitrypsin polymer specific $mAb_{2C1}$ antibody (HM2289, Hycult Biotechnology) was used for Native-PAGE detection of polymers.

**Z-A1AT polymer immunohistochemistry**

Immunohistochemistry staining was performed on liver tissue sections from subjects with A1AT-deficiency, adjacent to those on which LCM was previously performed. 5 µm thick sections were cut from Paxgene-fixed paraffin embedded tissue blocks and mounted on Superfrost Plus glass microscope slides. Sections were dewaxed by sequential immersion in xylene 2x for 2 min, 100% ethanol 2x for 2 min, 70% ethanol for 1 min and deionized water for 1 min. Endogenous peroxidase was blocked by soaking slides in methanol with 0.3% hydrogen peroxide solution for 30 min. After washing the slides in TBS, off-target secondary antibody was blocked by soaking the tissue sections in 3% normal horse serum (VECTASTAIN Elite ABC-HRP Mouse IgG

Kit, Vector Laboratories) in TBS for 1 hour. Tissue sections were washed in TBS and incubated overnight in 1:50 dilution of the mouse monoclonal 2C1 anti-Z-A1AT polymer-specific antibody (Hycult Biotech, Cat #HM2289). The next day, tissue sections were incubated with horseradish peroxidase (HRP)-conjugated horse anti-mouse IgG secondary antibody (1:200, VECTASTAIN Elite ABC-HRP Mouse IgG Kit, Vector Laboratories) for 1 hour. After removal of unbound antibodies, HRP activity was developed with diaminobenzene. The sections were then counterstained with Mayer's Hematoxylin, rinsed, rehydrated, mounted and imaged using a NanoZoomer 2.0-HT slide scanner (Hamamatsu Photonics, Hamamatsu, Japan).

2C1 staining intensity from regions of interest (ROI) was independently scored (0-3) by 4 independent scorers (VK, JC, SJM, TC) who were blinded to the genotype of the ROI and then a mean of these scores reported as the 2C1 staining intensity score. ROIs were grouped based on *SERPINA1* genotype: non-truncating variants included: L12delFS; F47InsFS; L124delFS; L144delFS; Q180delFS; K192insFS; V205delFS; Q236delFS; L397delIF. Truncating variants included: K367*; G373delFS; G373insFS M382delFS; S383insFS; P386insFS; E387insFS; E387*; F396delFS.

**HaloTag pulse-chase of A1AT**

CHO parental cells were transfected using Invitrogen™ Neon™ Transfection System with 5 µg of DNA encoding HaloTag-A1AT (M, Z, Z-K367*, Z-E387* variants) per $5x10^5$ cells using 3 electric pulses at 1400 V and 10 ms width. Cells were seeded at a density of $8.68x10^3$ cells/$cm^2$ in 6-well cell culture plates (Greiner Bio-One, UK). Cells were labelled with PA-JF646 photoactivatable ligand after 42 hours and chased for a further 24 hours. Cells were harvested at 0 hours, 1.5 hours, 3 hours, 6 hours, 12 hours and

24 hours as previously described. Cell lysates were sonicated in a water bath sonicator for 30 minutes at 4°C (whole cell lysate). 60µg of protein lysates from the first timepoint and an equivalent volume of protein lysates from subsequent timepoints were prepared in SDS-loading buffer as previously described for SDS-PAGE analysis. Gel fluorescence was detected using a Li-Cor CLx scanner.

**HaloTag pulse-chase of Z-K367* with inhibitors of degradation**

CHO parental cells were transfected using Invitrogen™ Neon™ Transfection System with 5 µg of DNA encoding HaloTag-Z-K367* per $5x10^5$ cells by Invitrogen™ Neon™ Transfection System using 3 electric pulses at 1400 V and 10 ms width. Cells were seeded at a density of $8.33x10^3$ cells/$cm^2$ in 6-well cell culture plates (Greiner Bio-One, UK). Cells were labelled with PA-JF650 photoactivatable ligand after 42 hours and substituted with media containing 5 µM lactacystin, 100 nM bafilomycin or both and chased for a further 6 hours. Cells were harvested at 0 hours, 1.5 hours and 3 hours and 6 hours. Cell lysates were centrifuged at 16,100xg for 10 minutes at 4°C in a bench-top centrifuge and supernatant was separated from pelleted material as previously described. 50µg of protein lysates from the first timepoint and equivalent volumes of proteins lysates from subsequent timepoints were prepared in SDS-loading buffer for SDS-PAGE analysis as previously described. Gel fluorescence was detected using a Li-Cor CLx scanner.

**Halo-Link pulldown of A1AT**

CHO Tet-ON cells with inducible expression of untagged Z-A1AT were seeded at a density of $2.08x10^6$cells/$cm^2$ in 6-well cell culture plates (Greiner Bio-One, UK). Transfection was performed as described above for parental CHO with HaloTag-$\alpha_1$-

antitrypsin (M, Z, Z-K367*, Z-E387* variants). Cells were next treated with 1μg/mL doxycycline for the induction of Z-A1AT expression for 48 hours. Cells were harvested by washing twice with PBS on ice prior to applying 200μL of ice-cold lysis buffer (50mM Tris-HCl pH7.4, 150mM NaCl, 1% Triton) containing 1X protease inhibitor (G6521, Promega). Cell homogenates were collected and spun at 14,000xg for 5 minutes at 4°C in a bench-top centrifuge. The cell lysate supernatant was used for the subsequent steps of the HaloLink pulldown analysis. Equal amounts of protein lysates were incubated with Magne HaloTag beads (#G7281, Promega) overnight at 4°C in a rotor. Beads were washed 4 times with lysis buffer and protein interactors of HaloTag fusion proteins were liberated with 2X SDS-PAGE sample buffer (312.5mM Tris-HCl pH6.8, 50% v/v glycerol, 10% w/v SDS, 0.05% w/v bromophenol blue, 50mM DTT) by heating at 75°C for 10 minutes and separation from HaloTag bead. SDS-PAGE was performed on eluted material and 80μg of input, followed by western blotting using the method described above.

**HaloTag pulse-chase of A1AT**

CHO parental cells were transfected using Invitrogen™ Neon™ Transfection System with 5 μg of DNA encoding HaloTag-A1AT (M, Z, Z-K367*, Z-E387* variants) per $5x10^5$ cells using 3 electric pulses at 1400 V and 10 ms width. Cells were seeded at a density of $8.68x10^3$ cells/$cm^2$ in 6-well cell culture plates (Greiner Bio-One, UK). Cells were labelled with PA-JF646 photoactivatable ligand after 42 hours and chased for a further 24 hours. Cells were harvested at 0 hours, 1.5 hours, 3 hours, 6 hours, 12 hours and 24 hours as previously described. Cell lysates were sonicated in a water bath sonicator for 30 minutes at 4°C (whole cell lysate). 60μg of protein lysates from the first timepoint and an equivalent volume of protein lysates from subsequent timepoints were prepared in SDS-loading buffer as previously described for SDS-PAGE analysis. Gel fluorescence was detected using a Li-Cor CLx scanner.

**HaloTag pulse-chase of K367***

CHO parental cells were transfected using Invitrogen™ Neon™ Transfection System with 5 µg of DNA encoding HaloTag-Z-K367* per $5x10^5$ cells by Invitrogen™ Neon™ Transfection System using 3 electric pulses at 1400 V and 10 ms width. Cells were seeded at a density of $8.33x10^3$ cells/cm$^2$ in 6-well cell culture plates (Greiner Bio-One, UK). Cells were labelled with PA-JF650 photoactivatable ligand after 42 hours and substituted with media containing 5 µM lactacystin, 100 nM bafilomycin or both and chased for a further 6 hours. Cells were harvested at 0 hours, 1.5 hours and 3 hours and 6 hours as previously described. Cell lysates were centrifuged at 16,100xg for 10 minutes at 4°C in a bench-top centrifuge and supernatant was separated from pelleted material as previously described. 50µg of protein lysates from the first timepoint and equivalent volumes of proteins lysates from subsequent timepoints were prepared in SDS-loading buffer for SDS-PAGE analysis as previously described. Gel fluorescence was detected using a Li-Cor CLx scanner.

**Statistics**

Statistical analyses were performed using R as indicated in the methods or using Graphpad Prism 9. The statistical tests used and the P values are described in figure legends. No data were excluded from the analyses. The experiments were not randomized and the investigators, apart from the pathologist, were not blinded to allocation during the experiments and outcome assessment.

**Data availability**

Sequencing data will be deposited at the European Genome-Phenome Archive upon publication.

#### References


1. Girelli, D. *et al.* Hemochromatosis classification: update and recommendations by the BIOIRON Society. *Blood* **139**, 3018–3029 (2021).

2. Ng, S. W. K. *et al.* Convergent somatic mutations in metabolism genes in chronic liver disease. *Nature* **598**, 473–478 (2021).

3. Brunner, S. F. *et al.* Somatic mutations and clonal dynamics in healthy and cirrhotic human liver. *Nature* **574**, 538–542 (2019).

4. Kleiner, D. E. *et al.* Design and validation of a histological scoring system for nonalcoholic fatty liver disease. *Hepatology* **41**, 1313–1321 (2005).

5. Scheuer, P. J., Williams, R. & Muir, A. R. Hepatic pathology in relatives of patients with haemochromatosis. *J. Pathol. Bacteriol.* **84**, 53–64 (1962).

6. Dawwas, M. F., Davies, S. E., Griffiths, W. J. H., Lomas, D. A. & Alexander, G. J. Prevalence and Risk Factors for Liver Involvement in Individuals with PiZZ-related Lung Disease. *Am. J. Respir. Crit. Care Med.* **187**, 502–508 (2013).

7. Ishak, K. *et al.* Histological grading and staging of chronic hepatitis. *Journal of Hepatology* **22**, 696–699 (1995).

8. Ellis, P. *et al.* Reliable detection of somatic mutations in solid tissues by laser-capture microdissection and low-input DNA sequencing. *Nat Protoc* 1–31 (2020) doi:10.1038/s41596-020-00437-6.

9. Ellis, P. *et al.* Reliable detection of somatic mutations in solid tissues by laser-capture microdissection and low-input DNA sequencing. *Nat. Protoc.* **16**, 841–871 (2021).

10. Li, H. & Durbin, R. Fast and accurate short read alignment with Burrows–Wheeler transform. *Bioinformatics* **25**, 1754–1760 (2009).

11. Tischler, G. & Leonard, S. biobambam: tools for read pair collation based algorithms on BAM files. *Source Code Biol. Med.* **9**, 13–13 (2014).

12. Jun, G. *et al.* Detecting and Estimating Contamination of Human DNA Samples in Sequencing and Array-Based Genotype Data. *Am. J. Hum. Genet.* **91**, 839–848 (2012).

13. Jones, D. *et al.* cgpCaVEManWrapper: Simple Execution of CaVEMan in Order to Detect Somatic Single Nucleotide Variants in NGS Data. *Curr. Protoc. Bioinform.* **56**, 15.10.1-15.10.18 (2016).

14. Raine, K. M. *et al.* cgpPindel: Identifying Somatically Acquired Insertion and Deletion Events from Paired End Sequencing. *Curr. Protoc. Bioinform.* **52**, 15.7.1-15.7.12 (2015).

15. Coorens, T. H. H. *et al.* Extensive phylogenies of human development inferred from somatic mutations. *Nature* **597**, 387–392 (2021).

16. Cameron, D. L. *et al.* GRIDSS: sensitive and specific genomic rearrangement detection using positional de Bruijn graph assembly. *Genome Res.* **27**, 2050–2060 (2017).

17. Loo, P. V. *et al.* Allele-specific copy number analysis of tumors. *Proc. Natl. Acad. Sci.* **107**, 16910–16915 (2010).

18. Raine, K. M. *et al.* ascatNgs: Identifying Somatically Acquired Copy-Number Alterations from Whole-Genome Sequencing Data. *Curr. Protoc. Bioinform.* **56**, 15.9.1-15.9.17 (2016).

19. Alexandrov, L. B. *et al.* The repertoire of mutational signatures in human cancer. *Nature* **578**, 94–101 (2020).

20. Nik-Zainal, S. *et al.* The Life History of 21 Breast Cancers. *Cell* **149**, 994–1007 (2012).

21. Martincorena, I. *et al.* Universal Patterns of Selection in Cancer and Somatic Tissues. *Cell* **171**, 1029-1041.e21 (2017).

22. Elliott, P. R., Pei, X. Y., Dafforn, T. R. & Lomas, D. A. Topography of a 2.0 Å structure of α1-antitrypsin reveals targets for rational drug design to prevent conformational disease. *Protein Sci.* **9**, 1274–1281 (2000).

23. Yamasaki, M., Sendall, T. J., Pearce, M. C., Whisstock, J. C. & Huntington, J. A. Molecular basis of α1-antitrypsin deficiency revealed by the structure of a domain-swapped trimer. *EMBO Rep.* **12**, 1011–1017 (2011).

24. Abramson, J. *et al.* Accurate structure prediction of biomolecular interactions with AlphaFold 3. *Nature* **630**, 493–500 (2024).

25. Dickens, J. A. *et al.* The endoplasmic reticulum remains functionally connected by vesicular transport after its fragmentation in cells expressing Z-α1-antitrypsin. *FASEB J.* **30**, 4083–4097 (2016).

26. Chambers, J. E. *et al.* Z-α1-antitrypsin polymers impose molecular filtration in the endoplasmic reticulum after undergoing phase transition to a solid state. *Sci. Adv.* **8**, eabm2094 (2022).

27. Grimm, J. B. *et al.* Bright photoactivatable fluorophores for single-molecule imaging. *Nat. Methods* **13**, 985–988 (2016).

28. Tinevez, J.-Y. *et al.* TrackMate: An open and extensible platform for single-particle tracking. *Methods* **115**, 80–90 (2017).
